# IsoProt: A complete and reproducible workflow to analyse iTRAQ/TMT experiments

**DOI:** 10.1101/446070

**Authors:** Johannes Griss, Goran Vinterhalter, Veit Schwämmle

## Abstract

Reproducibility has become a major concern in biomedical research. In proteomics, bioinformatic workflows can quickly consist of multiple software tools each with its own set of parameters. Their usage involves the definition of often hundreds of parameters as well as data operations to ensure tool interoperability. Hence a manuscript’s methods section is often insufficient to completely describe and reproduce a data analysis workflow. Here we present IsoProt: A complete and reproducible bioinformatic workflow deployed on a portable container environment to analyse data from isobarically-labeled, quantitative proteomics experiments. The workflow uses only open source tools and provides a user-friendly and interactive browser interface to configure and execute the different operations. Once the workflow is executed, the results including the R code to perform statistical analyses can be downloaded as an HTML or PDF document providing a complete record of the performed analyses. IsoProt therefore represents a reproducible bioinformatics workflow that will yield identical results on any computer platform.

## Introduction

Lack of reproducibility in general, and in bioinformatics workflows specifically is a growing concern^1^. Bioinformatic workflows in proteomics experiments often consist of multiple software tools, each with its own set of parameters. Seemingly small changes to a workflow, such as using different normalisation method details can have dramatic effects on the final result. Due to the many steps and settings that make a complex workflow, it is often impossible to fully document it in a research paper’s methods section. Additionally, finding and using the exact same software versions later on often represents a major obstacle when replicating bioinformatic analyses. Older versions may no longer be compatible with the available operating system or are just altogether unavailable. Therefore, fully reproducible workflows should not only record the exact software versions and parameters, but also preserve specific software versions and ensure that they will produce the same results in different computing environments.

Several projects exist to create reproducible bioinformatic workflows. Biocontainers^2^ provides Docker containers to make bioinformatic tools available in a standardized way. Docker containers are lightweight virtual machines that, in case of Biocontainers, ensure, that a given software version performs identically on any operating system supported by Docker. Therefore, users do not have to install any software but only download the respective container. Galaxy^3^ is a web-based platform for biomedical research focused on genomics. It contains thousands of tools which can be joined together to create workflows and also supports tools for proteomics analyses. KNIME (http://www.knime.com, KNIME AG) is another workflow software focused on data analysis in general. All OpenMS^4^ nodes were recently integrated in KNIME making it possible to build complete proteomics workflows with it. ProteomeDiscoverer (Thermo Fisher) is also a workflow system but specifically targeting proteomics data analysis. Several academic research groups^4–6^ are contributing to ProteomeDiscoverer making it usable for a wide variety of proteomics workflows. Finally, to a certain extent MaxQuant^7^ with Perseus^8^ allow the user to create a complete analysis workflow in a single software.

Nevertheless, all of these existing solutions have shortcomings that prevent the creation of complete, reproducible workflows. Biocontainers is a platform to supply bioinformatic tools in a standardized fashion but has no functionality to combine these tools into workflows. KNIME and Galaxy are very powerful analysis platforms that can be adapted to a wide variety of data analysis problems. This functionality comes at the cost of high complexity and many non-expert users will find it difficult to adopt Galaxy and KNIME for their needs. Additionally, both KNIME and Galaxy do not contain methods to take a snapshot of the external tools used to actually process the data. ProteomeDiscoverer also depends on external nodes. Therefore, to fully replicate an existing workflow the user again has to take care of locating and installing the exact same versions of these nodes. Moreover, new ProteomeDiscoverer versions generally come with significant changes which requires nodes to be specifically developed for a given version. Nodes developed for one version of ProteomeDiscoverer are generally incompatible with newer ones. Therefore, none of these existing solutions fulfill all requirements of a completely reproducible workflow environment.

Isobaric labelling has become one of the most common methods for quantitative mass spectrometry based proteomics experiments. A major advantage is that it allows researchers to multiplex samples and thereby reduce instrument runtime and eliminate variability caused by the mass spectrometer itself. The two methods currently available for these experiments Tandem Mass Tag (TMT, Proteome Science) and Multiplexed Isobaric Tagging Technology for Relative Quantitation (iTRAQ^9^) basically only differ in the reporter masses they generate but do not require dedicated software tools.

Even though isobaric labelling has become a standard method in many laboratories, dedicated, easy-to-use software solution to analyse these data are still rare. This is particularly problematic when dealing with more complex experimental designs that include multiple runs on the mass spectrometer, such as multiple instances of differently labeled multiplexed samples. Therefore, many research groups rely on unpublished in-house scripts to process their experiments which greatly hampers reproducibility.

In an effort to simplify proteomics data analysis and provide fully reproducible data analysis workflows we launched the ProtProtocols project (https://protprotocols.github.io) under the umbrella of the European Bioinformatics Community (EuBIC)^10^. Based on the Biocontainers project^2^ the protocols are shipped in containerized Docker images that include all necessary software tools. Docker containers are lightweight virtual machines that encapsulate all the software required for the protocol to run. This ensures that the version of all used software is linked to the protocol version and the user does not have to worry about installing any separate tools. Hence, 100% reproducibility can be achieved by using the same protocol version on any computer with a Docker environment.

Here, we present IsoProt which serves as a blueprint for the ProtProtocol concept. IsoProt is designed for the analysis of isobarically labelled experiments which is one of the most commonly used methods for high-throughput proteomics. Next to a user-friendly web interface, IsoProt provides accurate statistical analyses for a wide range of common experimental designs.

## Experimental Procedures

### Software layout and implementation

#### General implementation

All software was installed in a Docker image to ensure full reproducibility on each computer system supported by Docker. To simplify the installation and usage of our protocols we created the free, open-source “ProtProtocol docker-launcher” (https://github.com/ProtProtocols/docker-launcher). It provides an easy-to-use graphical user interface that can automatically install the protocol (once Docker is installed) and launch the image. As it is written in Java it supports the major operating systems Windows, Mac OSX, and Linux. Therefore, many technical difficulties surrounding the use of Docker are hidden from the user.

The complete protocol is run through a Jupyter notebook (http://jupyter.org) corresponding to one web page in the browser. All relevant parameters can be set through common graphical user elements created through Jupyter widgets. Therefore, the user interface is highly similar to most available search engines. The complete source code as well as additional documentation of the protocol is freely available through https://protprotocols.github.io.

#### Proteomics software

IsoProt handles the entire analysis pipeline from mass spectra given as peak lists to the set of differentially regulated proteins (Figure 1A). We used SearchGUI^11^ and PeptideShaker^12^ to perform peptide identification and validation, with MS-GF+^13^ as database search engine. Proteins are summarized and quantified by R scripts based on the MSnBase R library^14^. R scripts furthermore generate figures for quality control and perform statistical tests (LIMMA library^15^) according to the experimental design.

**Figure 1:**
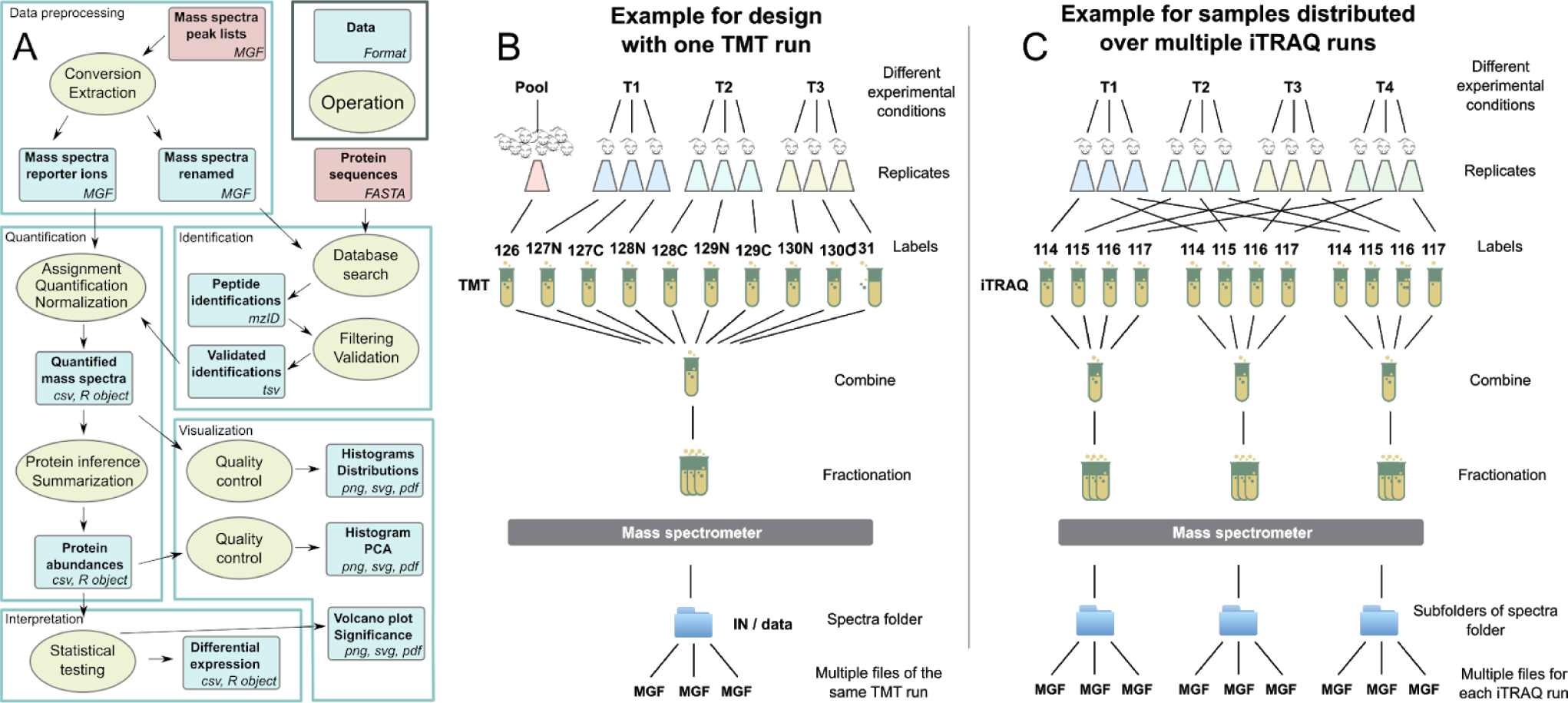
A Scheme of the entire workflow including operations (ellipsoids) and data given by type and format (squares). The annotation form and terms of the workflow follow to large extent the EDAM ontology^16^. B+C Experimental designs and organization in folder structure for analysis in IsoProt.

### Input files and parameters

#### Input files

The only files required for the analysis are mass spectra as peak lists (MGF format) and a FASTA file containing the protein sequences where we recommend the UniProt version of the FASTA format. Databases can already contain decoy sequences (following the SearchGUI instructions, http://compomics.github.io/projects/searchgui.html), otherwise the decoy database is created automatically. The files can be copied into the Docker file structure or directly mirrored onto the */data* folder as automatically done by our docker-launcher application.

#### Analysis parameters

All parameters required for the data analysis can be changed through a graphical user interface integrated into the Jupyter notebook. In the first section, the user has to set database search related parameters such as precursor and fragment ion tolerance, the FASTA sequence database to use, the labelling agent used, and the fixed and variable modifications to consider.

Based on the selected labelling method and detected folder structure, the interface to enter the experimental design is generated. The protocol currently supports two setups: 1) all MGF files are placed in the input directory and are part of the same (fractionated) run (Figure 1B) or 2) MGF files from different runs are organized by placing them in different subdirectories (Figure 1C). Next, the experimental design user interface allows the user to enter names for the sample groups (for example “treatment” and “control”), names for the samples (one name per channel and subdirectory) and assign each sample to one of the groups. Most importantly, the protocol supports up to 20 sample groups and can thereby model complex experimental designs.

Finally, the user is asked to enter parameters related to the analysis of the quantitative data. Once all required information is entered, the search and analysis is directly controlled through buttons in the user interface.

### Output files and quality control

IsoProt provides figures and tables for the different steps of the analysis including peptide identifications, quantitative values of peptide-spectrum matches (PSMs) and proteins as well as a table for the statistical results from the significance analysis. Visual measures for quality control were implemented as R scripts and include total intensities of the reporter ion channels for each sample, violin plots at different stages of the analysis, principal component analysis and volcano plots (Figure 2).

**Figure 2:**
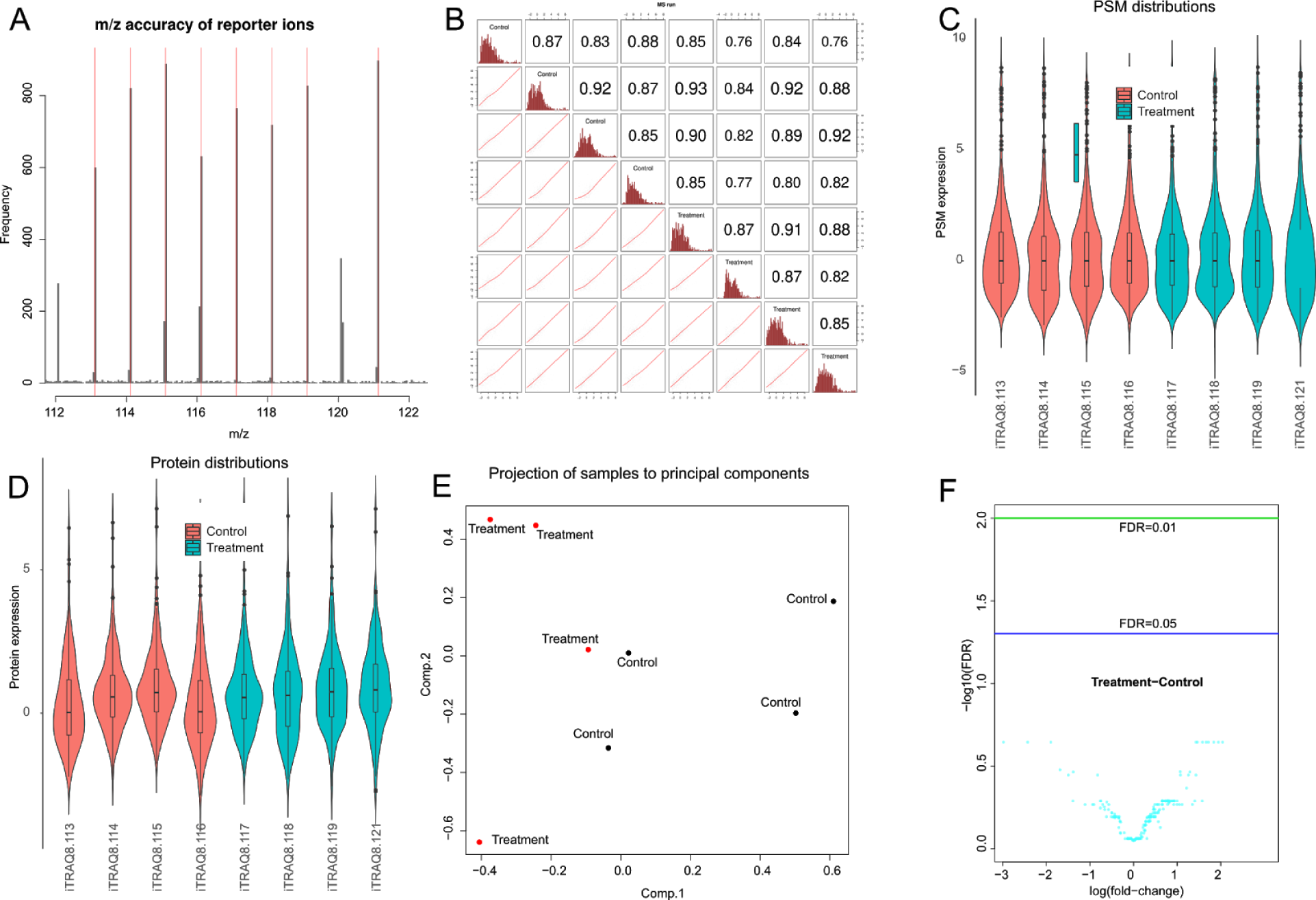
Examples of the visualisation and diagnostic plots created by IsoProt based on the shipped example data. A) The mass accuracy of all reporter ions is presented as a histogram. B) Correlation of reporter intensities for all channels. C) Distribution of estimated abundances on the spectrum level for all channels. D) Distribution of protein abundances for all samples. E) Principal component analysis of all samples based on the aggregated data highlighting the treatment groups. F) Volcano plot for a quick visualization of quantitative data and statistical results.

### Test data sets

To evaluate the performance of our analysis workflow we processed the data from three publically available datasets. We downloaded the respective RAW files from PRIDE Archive^17^ and converted them into the MGF file format using ProteoWizard’s msconvert tool^18^ when no MGF peak list files were available.

#### Benchmark dataset

D’Angelo *et al*. recently published a TMT benchmark dataset containing an experiment where 12 human proteins were spiked into an *E*. *Coli* background^19^ using various concentrations (PRIDE Archive identifier PXD005486). D’Angelo *et al*. used this dataset to assess the number of proteins that were incorrectly identified as being regulated. As every protein was added using varying concentrations amongst the samples, a standard statistical analysis of the spiked-in proteins was not possible. Therefore, our analysis focuses on the accuracy of the derived quantitative estimates for the spiked proteins and the (unchanged) background *E*. *Coli* proteins.

The complete analysis was performed using IsoProt version 0.2. Spectra were identified using MSGF+^13^ through SearchGui version 3.3.3^11^. The precursor tolerance was set to 20 ppm and the fragment tolerance to 0.03 Daltons. 1 missed cleavages were allowed. Carbamidomethylation and TMT 10-plex of K,TMT 10-plex of peptide N-term were set as fixed modifications. Oxidation of M was set as variable modification. PSMs were filtered at a target false discovery rate (FDR) of 0.01 using the target-decoy approach. UniProt E. Coli sequences (version August 2018) and the spiked human protein sequences, also from UniProt, were used for spectra identification.

Quantitative analysis was done using the R Bioconductor package MSnbase version 2.7.1^14^. Protein summarization was performed using the ‘medpolish’ method as implemented by MSnbase. Modified peptides were not used for quantitation. Only proteins with at least 2 identified peptides were accepted for further analysis. Differential expression was assessed using the R Bioconductor package limma version 3.34^15^.

#### Cerebral malaria pathogenesis

The study uses TMT6 labeling to compare mouse blood with different stages of cerebral malaria (d3, ECM) to non-infected mice (NI)^20^. Four replicates of each of the three sample types were arranged in TMT6 sets and run separately. Peak list data files (MGF file format) were downloaded from PRIDE (PXD003772).

The analysis was again performed using IsoProt version 0.2 (see above) with the precursor tolerance set to 10 ppm and the fragment tolerance to 0.05 Daltons. 1 missed cleavage was allowed. Carbamidomethylation and TMT 6-plex of K,TMT 6-plex of peptide N-term were set as fixed modifications. Oxidation of M was set as variable modification. PSMs were filtered at a target FDR of 0.01 using the target-decoy approach. SwissProt sequences from mouse (January 2018) were used for spectra identification. Only proteins with at least 2 identified peptides were accepted for further analysis.

#### N*on-muscle invasive and muscle-invasive bladder cancer*

The study compares tumor tissue samples from non-muscle invasive and muscle-invasive bladder cancer^21^. MGF files were downloaded from PRIDE Archive (PXD002170). The analysis was again performed using IsoProt version 0.2 (see above) with the precursor tolerance set to 10 ppm and the fragment tolerance to 0.05 Daltons. 1 missed cleavage was allowed. Carbamidomethylation and iTRAQ 8-plex of K,iTRAQ 8-plex of Y,iTRAQ 8-plex of peptide N-term were set as fixed modifications. Oxidation of M was set as variable modification. PSMs were filtered at a target FDR of 0.01 using the target-decoy approach. Sequences from SwissProt sequences from human (January 2017) were used for spectra identification. Only proteins with at least 2 identified peptides were accepted for further analysis.

## Results

IsoProt allows users running the full data analysis of iTRAQ/TMT experiments in a straight-forward and reproducible way. The protocol supports different experimental designs including multiple runs on the mass spectrometer and differently labeled multiple samples. Additionally, the open layout of the protocol allows complex adjustments and modifications at all stages of the workflow.

### A fully reproducible environment

The protocol can be run on any computer with a functional Docker environment, by just downloading and running the available Docker image. This is fully automated through our “ProtProtocol docker-launcher” tool (https://github.com/ProtProtocols/docker-launcher). Hence, the protocol avoids all possible platform- and operating system-specific installation issues and provides identical results independent of operating system, its configuration and computer hardware.

Every IsoProt release has a stable version number that points to a specific docker image. Therefore, by citing the used IsoProt version number it will always be possible to exactly restore the used analysis environment - including the versions of all used software tools. Once the protocol has been executed, it is possible to save it, including all generated figures, as a standard HTML or pdf page. Therefore, the complete analysis workflow can be easily made available, for example at the time of review, and be viewed with a standard web browser. For an overview of the visualizations, see Figure 2.

### Simple example workflow

IsoProt can be tested using an example data set which is small enough to run in under ten minutes on a standard computer. The data set is part of the IsoProt Docker image and necessary parameters settings are preloaded when starting IsoProt. The database search via SearchGUI and validation via PeptideShaker result in a tab-delimited file containing detailed information of all PSMs. Search and output parameters are automatically saved for future reference. Additionally, a “methods” section is generated that can be included in a manuscript and describes all used settings. Each spectra file is processed separately to match and quantify PSMs that passed the identification FDR (default 0.01). The mass distribution of all matched fragment ions allows control for critical channels with inefficient labeling (Figure 2A). All PSM quantifications are saved in a separate file (AllQuantPSMs.csv).

The output of all files of each run on the mass spectrometer are merged, normalized and visualized for quality control. Violin plots of normalized PSM intensities compare the intensity distributions (Figure 2C). Channels with different distributions can identify problematic samples or changes within the entire proteome. Six different histograms counting PSM, peptide, protein and protein group numbers allows determining protein coverage and uniqueness by the available mass spectra. Similarity between samples is assessed through scatter plots comparing all quantified spectra from all ion channels (Figure 2B).

Using the default parameters, the PSMs are summarized to proteins using median summarization after outlier removal requiring a minimum of 1 PSM per protein. Other methods such as iPQF<u>^22^</u> are available. A violin plot of protein ratios versus mean of all channels shows whether the analysed samples exhibit similar distributions on the protein level (Figure 2D).

Quantifications from different runs (only one in the example) are merged and submitted to a principal component analysis (Figure 2E). With the scoring plot, one can look for similarity within replicates compared to different experimental conditions. Completely mixed samples are unlikely to provide differentially regulated proteins.

The example set quantified a total of 221 protein groups. LIMMA statistical tests did not find any regulated proteins with FDR < 0.05 which is in agreement with the original results. p-values and false discovery rates (p-values corrected for multiple testing) are visualized in histograms, volcano plots (Figure 2F) and a figure counting the number of differentially regulated proteins over a range of FDRs. The latter can be used to identify a suitable combination of the confidence threshold and the number of significant proteins. It is advised to keep FDR<0.1 as the number of false positives becomes critically high otherwise.

### Performance tests by reprocessing public data

#### Benchmark dataset

D’Angelo *et al*. performed a comparison of different approaches to analyse TMT datasets^19^. In their first dataset the authors spiked different concentrations of 12 human proteins into an *E*. *Coli* background. They used this dataset to assess the type-I error as the number of false positive proteins. Similar to the original study, we assigned the first five channels to one treatment group, and the second five channels to the second group. As expected, no proteins were identified as being significantly regulated. The estimated log-fold changes of the *E*. *Coli* background proteins were all close to 0 (Figure 3A).

**Figure 3:**
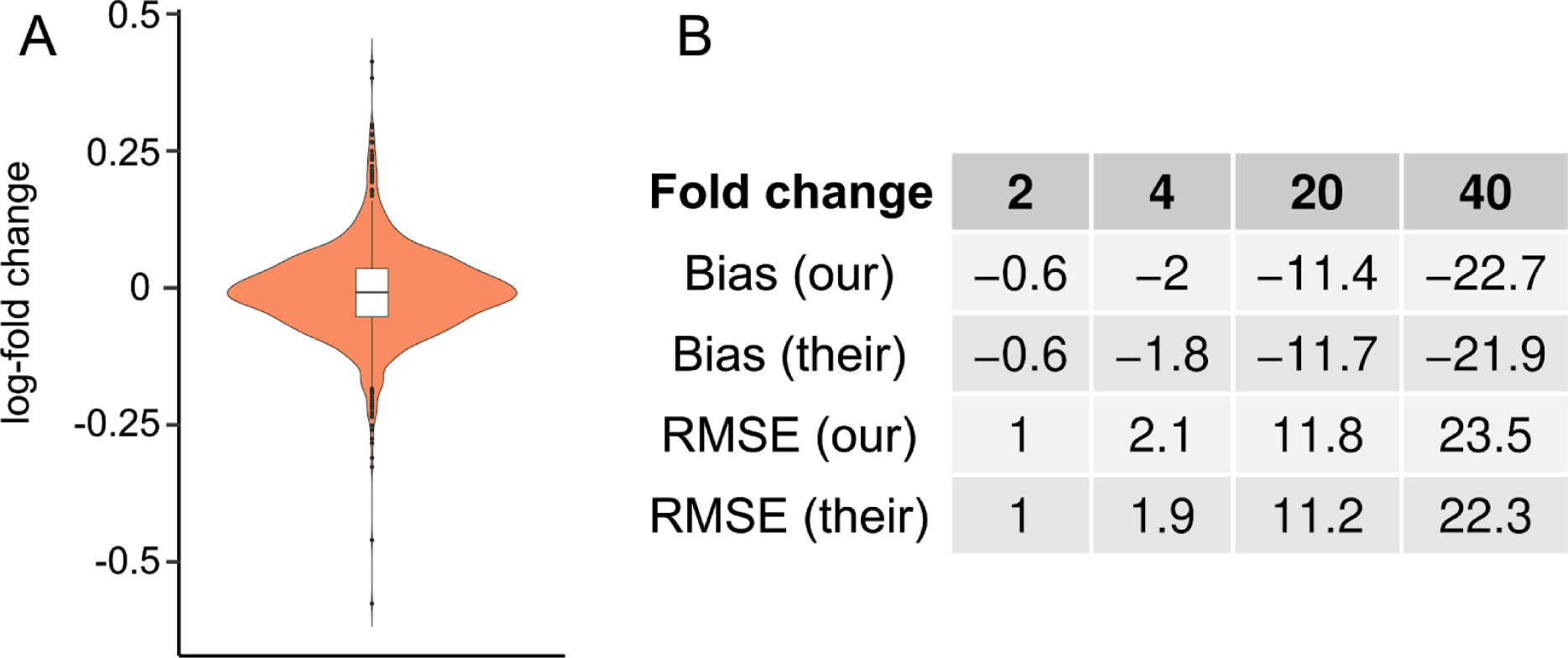
A Log-fold changes of the E. Coli background proteins. This represents the expression of background proteins based on a comparison of the first five channels against the other five ones similar to the approach by D’Angelo et al. As expected, the estimated log-fold changes are all closely centered around 0. B Observed bias and RMSE of estimated fold-changes of the D’Angelo et al. benchmark dataset from our pipeline and the best-performing pipeline published by the authors.

Proteins were spiked twice using the same concentration in different channels and only once for the two highest concentrations. Therefore, only a single, or two replicate measurements at maximum are available when comparing two concentrations. Since this setup prevents a standard statistical evaluation, we focused on the accuracy of the estimated fold changes using the same error measurements as in the original manuscript. Similarly, we assessed the accuracy of our fold change estimates using the bias and the root-mean-square error (RMSE). Across most spiked fold-changes, we observed a comparable bias and RMSE (Figure 3B).

For the highest spiked fold-change we observed slightly higher average error rates than D’Angelo *et al*. This is most likely caused by the fact the D’Angelo *et al*. imputed missing values by taking the lowest observed intensity of the given PSM across all samples. Thereby, missing values were automatically interpreted as very low expression. In our analysis, the measured abundance of the lowest concentrations was less stable. When we, for example, estimated the fold change of the two highest protein concentrations (also a fold change of 2), the bias is 0 with an RMSE of 0.2 improving the error rates dramatically.

While D’Angelo *et al*.’s imputation approach is valid if values can be expected to be missing not at random (ie. because of a concentration below the limit of detection) it is not valid for values missing (completely) at random (ie. because of inefficient labeling)^23^. Therefore, for the spiked proteins D’Angelo *et al*.’s approach should only have been applied to cases were the lowest amount of proteins were spiked. Since it is generally unknown why a value is missing in actual experiments, our pipeline is not using any imputation. The complete output of our pipeline can be found in Supplementary File 1.

#### Cerebral malaria pathogenesis

The authors investigated differences in the plasma proteome between healthy and malaria-infected mice (two stages). The available two TMT 6plex sets were considered to contain independent samples. IsoProt quantified more protein groups (324 versus 289) when requiring a minimum of 2 unique PSMs and an identification FDR < 1%. For the further comparison, we restricted the IsoProt output to the uniquely identified 214 proteins (no peptides shared with other proteins).

In the original study, statistical testing was carried out separately for the two TMT runs, yielding a total of 54 (more precisely 43 as 11 were detected in both runs) proteins found to be differentially regulated between *plasmodium berghei* ANKA (PbA)-infected (d8 post-infection, labeled ECM) and non-infected (labelled NI) mice (Mann-Whitney *U* test, p ≤ 0.001, no correction for multiple testing). We found a total of 41 differentially regulated proteins (FDR < 0.01) and an overlap of only 20 proteins with the original study.

Given the rather different procedure for statistical testing in the original analysis, we investigated proteins that were not determined as differentially regulated by either method. All but four proteins found differentially regulated in the original study were quantified by IsoProt and showed similar abundances in both analyses (Figure 4A). Proteins only found significantly changing in the original study were not found significant by IsoProt mostly due to low fold-changes in the quantitative analysis (Figure 4B).

**Figure 4:**
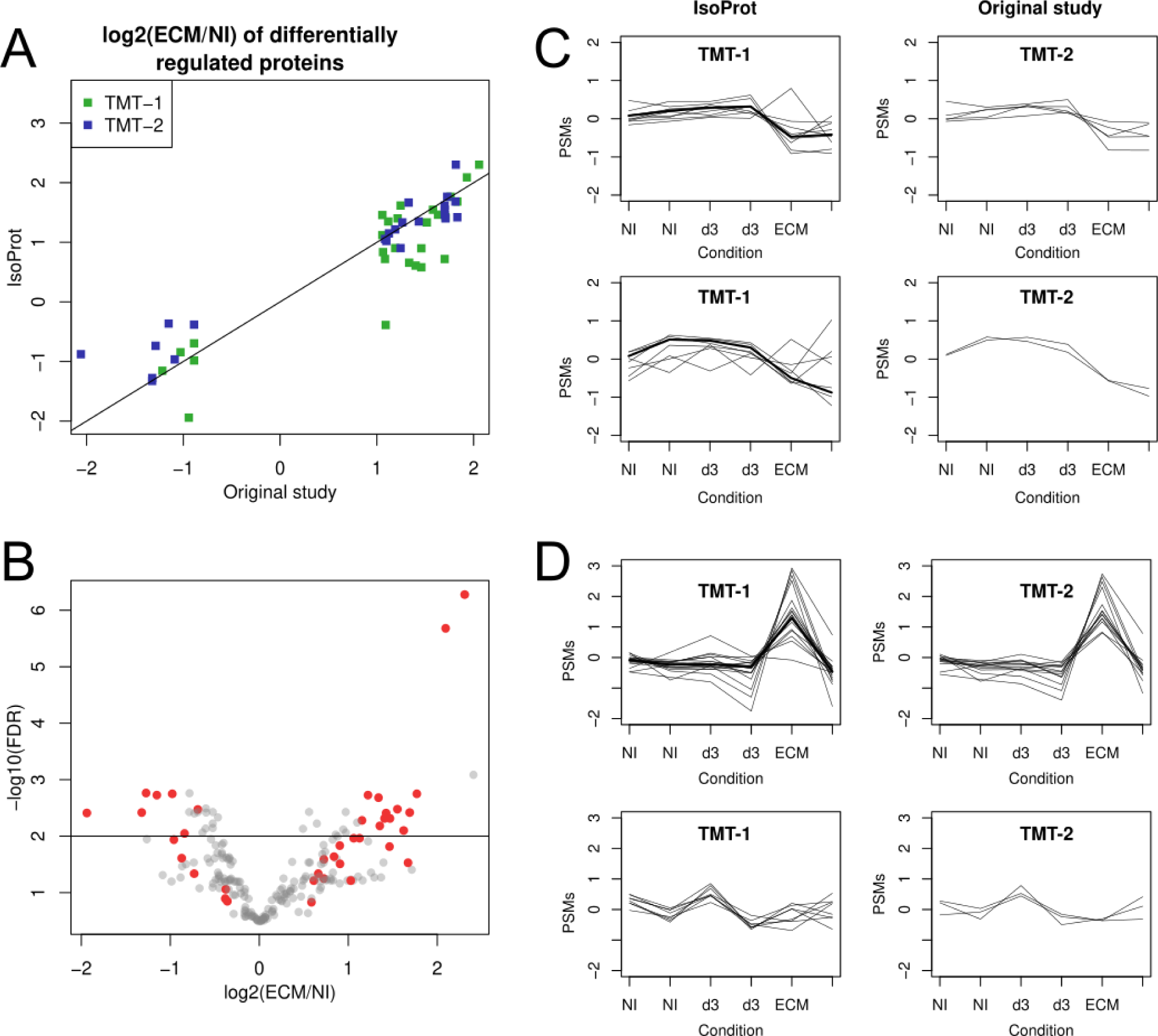
**A** Comparison of fold-changes of proteins differentially regulated in the original study and IsoProt results. **B** Volcano plot of IsoProt results. Proteins found differentially regulated in the original study were labeled red. **C** Abundance profiles of Retinol-binding protein 4 (Q00724). **D** Abundance profiles of disulfide-isomerase (P09103).

We investigated the two proteins that mostly differ between the 2 types of analyses. Retinol-binding protein 4 (Q00724) was the protein with the lowest FDR within the proteins found differentially regulated by IsoProt but not in the original study. Figure 4C shows PSM measurements for the 2 TMT runs of this protein (scaled for better comparison).

Summarized protein abundances (thick lines) by median summarization with outlier removal show that the PSMs of peptides with less differential behavior were removed. By merging the observation of the two TMT runs, IsoProt gains more statistical power and thus provides evidence for regulatory behavior of this protein.

On the other hand, protein Protein disulfide-isomerase (P09103) was the protein with the highest FDR (least significant) of proteins found significantly changing in the original study (TMT-1) but not by IsoProt (Figure 4D). Given only high abundances in one of the two ECM replicates in TMT-1, at least manual interpretation would discard this protein from being regulated (Figure 4D). The PSMs measured in the 2nd TMT-2 run confirm this observation. The complete output of our pipeline can be found in Supplementary File 2.

#### Non-muscle invasive and muscle-invasive bladder cancer

IsoProt quantified 1,145 protein groups when restricting to a minimum of 2 unique peptides and 1% FDR, compared to 1,092 in the original study (minimum of 2 peptides, Occam razor principle for peptide inference and 1% FDR). Both analyses had an overlap of 662 proteins.

We then compared the mean log-ratios between the two cancer subtypes (four replicates each). Despite only having different bioinformatics workflows, relatively large differences were observed between the estimated log-fold changes (Figure 5A, Pearson’s correlation of 0.78).

**Figure 5:**
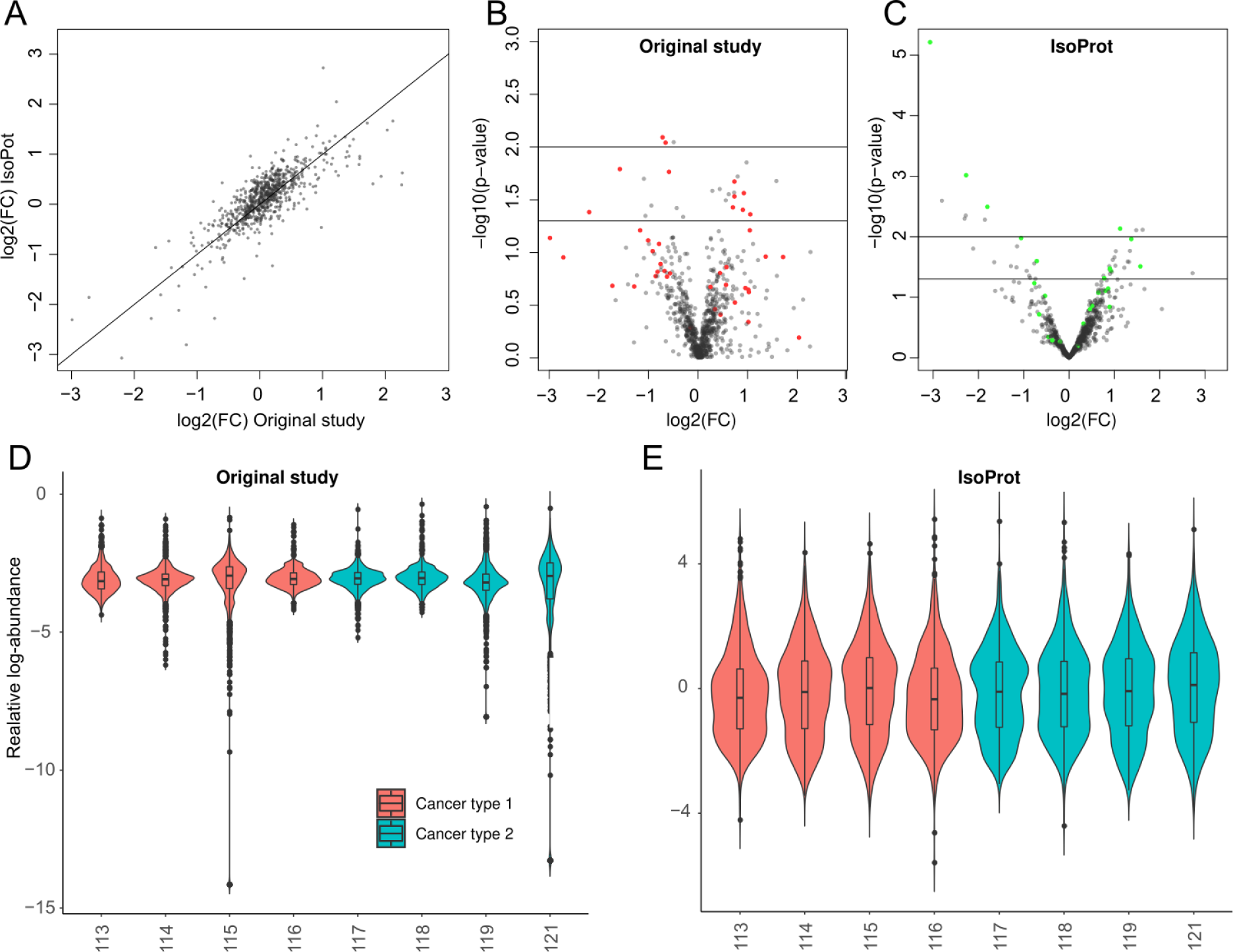
**A** Comparison of log-ratios between IsoProt output and original study. Pearson’s correlation between both quantification: 0.79. **B-C** Volcano plots for results from statistical testing in the original study (**B**) and in IsoProt (**C**). Colored points correspond to proteins with a p-value below 5% in the other study, respectively. **D-E** Distribution of relative protein abundances in original study (**D**) and IsoProt (**E**).

Statistical testing did show one differentially regulated protein (15-hydroxyprostaglandin dehydrogenase, FDR < 0.01) after correction for multiple testing which has not been carried out in the original study. When comparing uncorrected p-values, the majority of significant proteins were different between the two studies (Figure 5B and C, colored points indicate p<0.05 in the other respective study).

This striking difference in the statistical results can be explained by the distribution of protein abundances (Figure 5D and E). The authors of the original study normalized the ratios between cancer subtypes after protein summarization and averaging of replicates. The more common and in our opinion correct approach is to normalize the different channels (ie. individual samples) on the (measured) PSM or (aggregated) peptide level prior to the aggregated analysis of these measurements on the protein level and, most importantly, prior to merging any independent (i.e. replicate) measurements. Strong deviations of individual channels which are visible on the peptide level were thus discarded in the original study. The complete output of our pipeline can be found in Supplementary File 3.

## Discussion

IsoProt shows how the ProtProtocols framework can be used to create user-friendly, reproducible bioinformatic workflows. IsoProt makes it simple to include the complete bioinformatic data processing workflow as a supplementary file. Thereby, reviewers and other researchers can easily assess the used methods.

Encapsulating protocols into docker containers preserves the complete setup including all software versions which can be referenced through a single protocol version number. This allows anyone to replicate the results at any later stage without having to worry that older software might no longer work. Once a given version of the protocol is downloaded, users can be sure that it will behave in exactly the same way on all supported platforms.

The use of docker makes the protocol highly portable. Docker currently supports Windows, Linux and Mac OS making our protocol trully multiplatform. The fact that the protocol can be installed through a single command makes it trivial to move the setup from one machine to another. With our “ProtProtocol docker-launcher” tool the protocol can even be installed with the click of a single button. This should greatly reduce the effort in setting up a complex proteomics analysis environment. Unfortunately, Docker support for Windows is not yet fully stable. Therefore, several Windows users experienced issues when installing Docker which prevented them from using IsoProt. Even though this currently reduces the ease-of-use of ProtProtocols on Windows machines, we believe that this will quickly be improved since Microsoft recently became an official partner of Docker^24^.

IsoProt’s performance was tested on three publicly available datasets. The results highlight that subtle differences in the data analysis can lead to considerable differences in the final results. Such differences can only be identified by reproducing the complete environment of the analysis workflow, something that is very difficult to realize when only relying on information from a scientific paper. Thus, more complete and easily readable information of the used workflow and its parameters, or even the entire computational environment, will considerably improve paper reviews as well as reproducing and discussing results from already published studies. Such workflows will further increase quality and credibility of both scientific studies and the presenting journals. IsoProt enables users to easily provide such complete information on their analysis. Our approach facilitates comparison with other data analysis pipelines or testing of robustness to parameter changes with minimal efforts requiring only peak list files, their relation to the experimental design and main parameters for identification and quantification.

All of these developments are available as free and open-source software. Thereby, we encourage other researchers to use the ProtProtocol infrastructure as starting point to develop their own analysis workflows and make them available to the community. All our tools are modularized and prepared to support and simplify such external developments. Since Docker has become an industry standard for containerized applications long-term support seems to be guaranteed for these developments.

In summary, we developed a user-friendly environment for fully reproducible data analysis and exemplified its use through a complete workflow for the analysis of data from isobarically labelled mass spectrometry experiments.

## Acknowledgments

This project has received funding from the *European Research Council (ERC) under the European Union’s Horizon 2020 research and innovation programme* under grant agreement No 788042. JG additionally acknowledges financial support from the FWF-Austrian Science Fund [grant number P30325-B28]. VS acknowledges financial support from the Danish Research Council (DNRF82) and the EU ELIXIR consortium (Danish node). This project would not have been possible without support by the EuBIC initiative.

## Data Availability

The complete software presented in this manuscript is freely available under a permissive open-source license at https://protprotocols.github.io.

## Abbreviations

EuBIC: European Bioinformatics Community
FDR: False Discovery Rate
iTRAQ: Multiplexed Isobaric Tagging Technology for Relative Quantitation
PbA: *Plasmodium Berghei* ANKA
PSM: Peptide Spectrum Match
RMSE: Root-Mean-Square Error
TMT: Tandem Mass Tag

