## Supplementary File 1 for "IsoProt: A complete and reproducible workflow to analyse iTRAQ/TMT experiments"

Isobaric\_Workflow


### Protocol for analysis of labeled proteomics data¶

#### Introduction¶

This jupyter notebook contains a complete workflow to process and quantify TMT and iTRAQ labelled MS/MS data. The analysis ranges from peak lists to differentially regulated proteins. The protocol can be adapted to different experimental designs, different search parameters and several protein quantification methods.
The protocol is portable and reproducible, and should function on every computer that runs Docker.

#### Usage¶

**To run the example**, See below.

**To analyze your own data**

1. Convert raw MS data to MGF files (PeptideShaker currently requires all input files to be in the MGF format). We recommend ProteoWizard.
2. Use the "Home" screen of the Jupyter environment (normally at http://0.0.0.0:8888), navigate to the `IN` (delete iTRAQCancer.mgf before) or `data` directory and copy all MGF files there (using the `Upload` button)
3. Scroll down in this document until you reach the `Workflow` section to start processing your data.
4. All data files will be written to the folder **OUT**

**Resume a previous analysis**

The python scripts automatically detect if a previous' analysis results are present in the result directory ("OUT" in the directory where the "run" script was launched).

**Loading previous settings**

To reuse the settings of a previous analysis in a new project, follow these steps:

1. Create a new "OUT" folder in the directory where you want your new results to be placed
2. Copy the `protocol_parameters.json` (contains all search parameters) and the `exp_design.tsv` (contains the experimental design) files into the new OUT folder.

   - **WARNING:** The experimental design can only be re-used if the directory structure of the MGF files are exactly the same. This will probably not be the case. Therefore, in most cases we recommend to only copy the `protocol_parameters.json` file and re-eneter the experimental design.
3. Launch the docker container (using `run.sh` on Unix systems and `run.bat` on Windows) from the directory containing the "OUT" folder.

#### Maintainers¶

- Veit Schwämmle
- Johannes Griss
- Goran Vinterhalter

#### Software¶

Database searches are performed using searchGUI and the subsequent search results are filtered using PeptideShaker. Down-stream data processing in R used the Bioconductor libraries MSnbase and LIMMA amongst others.

bio.tools links for more detailed information about the used software:  
https://bio.tools/searchgui  
https://bio.tools/peptideshaker  
https://bio.tools/msnbase  
https://bio.tools/limma

#### Diagram¶

We used the EDAM ontology. Click on the links to get more information:  
Peptide data base search  
Target-decoy  
Labeled quantification  
Differential protein expression analysis  
Visualization

#### System requirements¶

Fill in the following items:
Required hard disk space for docker image, input and output files:
You will need space for your raw files and files from the down-stream analysis (mostly < 1 GB)

Required memory: Recommend min. 4 GB or RAM

Recommmended number of threads: 4-8

#### Example¶

The example data file is an extract of spectra from the iTRAQ 8-plex data in ref. https://doi.org/10.1371/journal.pone.0137048
The parameters of the example are preloaded. You can run it by going to the tab "Experimental design" below and press the "Save design" and "Run my search" buttons. The database search and the R analysis can take a few minutes.

### Workflow¶

In [2]:

```
from button_execute import ExecuteButton
# when clicking the button, the next 3 Jupyter notebook cells will be executed
ExecuteButton(button_text="Start Wokflow", n_next_cells=4)
```

In [4]:

```
%%javascript 
// disable scrolling for all cells
requirejs("notebook/js/outputarea").OutputArea.prototype._should_scroll = function(lines) {return false;}
```

In [5]:

```
#%load_ext rpy2.ipython
get_ipython().run_line_magic('load_ext', 'rpy2.ipython')
```

##### Specify parameters for database search and evaluation of identified peptide-spectrum matches:¶

In [6]:

```
# This cell contains the complete code to
# 1.) Display the GUI for the user to enter the search parameters
# 2.) Launch the search based on these parameters
# 3.) Display a button once the search is complete to execute the subsequent R analysis code

from shutil import copyfile
from IPython.display import display

search_in = None

def complete_function():
    global search_in
    
    search_in = search_ui_out.copy()
    
    # remove the search_ui object
    if "search_ui" in search_in:
        del search_in["search_ui"]
    
    search_in["on_search_complete"] = search_complete
    %run "Scripts/search.ipy"
    
def search_complete():    
    # Enable the R analysis button
    btn_analysis.disabled = False
    btn_analysis.button_style = "success"
    print("Start the analysis by clicking the >>Run R Analysis<< button")
    
def update_search_parameters(arg=None):
    """
    Update all settings parameters
    """
    global search_ui_out, search_in
    search_ui = search_ui_out["search_ui"]
    
    if not search_ui:
        raise Exception("Failed to retrieve searchUI object")
        
    # update the current settings
    search_ui_settings = search_ui.get_settings_as_dict()
    
    for setting, value in search_ui_settings.items():
        search_in[setting] = value
        
    # save the settings again
    search_ui.save_config("/home/biodocker/OUT/protocol_parameters.json")
    
def on_change_input_dir(search_ui, spectra_dir):
    # test if it is our test directory
    test_file = os.path.join(spectra_dir, "iTRAQCancer.mgf")
    exp_design_file = os.path.join(spectra_dir, "exp_design_example.tsv")
    target_exp_design_file = os.path.join("OUT", "exp_design.tsv")
    
    if "biodocker/IN" in spectra_dir and os.path.isfile(test_file) and os.path.isfile(exp_design_file) and not os.path.isfile(target_exp_design_file):
        # copy the experimental design file to the default location
        copyfile(exp_design_file, target_exp_design_file)
        search_ui.load_exp_design()
        
# ----------- ENTRY POINT ------------
search_ui_in = {"on_complete_description": "Run my search", "on_complete_function": complete_function,
            "on_change_input_dir": on_change_input_dir}

%run "Scripts/search_ui.ipy"

# create the R analysis button
btn_analysis = ExecuteButton(button_text="Run R analysis", n_next_cells=13, disabled = True)
btn_analysis.on_click(update_search_parameters)
display(btn_analysis)

# Test if results are already present and show the R analysis button in this case
if os.path.isdir("OUT") and os.access("OUT", os.R_OK) and \
   os.path.isfile("OUT/protocol_parameters.json") and \
   os.path.isfile("OUT/exp_design.tsv"):
    # prepare the objects
    search_in = search_ui_out.copy()
    if "search_ui" in search_in:
       del(search_in["search_ui"])

    # Test whether only one directory or multiple directories exist
    if os.path.isfile("OUT/experiment1_test_1_Extended_PSM_Report.txt"):
        # replicate all global variables as if the search was run
        print("Previous results found in OUT.")
        search_out = {"result_files": ["OUT/experiment1_test_1_Extended_PSM_Report.txt"]}
        search_complete()
    else:    
        result_files = list()
        
        # Test every directory whether it contains result files
        for file in os.listdir("OUT"):
            file_path = os.path.join("OUT", file)
            if os.path.isdir(file_path) and os.access(file_path, os.R_OK):
                result_file_path = os.path.join(file_path, "experiment1_test_1_Extended_PSM_Report.txt")
                if os.path.isfile(result_file_path):
                    result_files.append(result_file_path)
        
        if len(result_files) > 0:
            print(str(len(result_files)) + " result files found.")
            search_out = {"result_files": result_files}
            search_complete()
```

```
Previous results found in OUT.
Start the analysis by clicking the >>Run R Analysis<< button
Saving parameters to /home/biodocker/OUT/protocol_parameters.json
```

#### Methods Sections¶

In [7]:

```
%run Scripts/methods_generator.ipy
```

##### Convert Search Settings into R Objects¶

These cells should not produce any output.

In [8]:

```
# Since py2ri cannot convert dict objects, simply save everything R needs as a JSON string

# remove the callback function first
search_in.pop("on_search_complete", None)

import json
search_in_string = json.dumps(search_in)
search_out_string = json.dumps(search_out)

import rpy2.robjects as ro
from IPython.display import display,HTML
import sys
```

In [9]:

```
%%R -i search_in_string,search_out_string -o samples
# Convert the JSON objects back into "natural" R objects
suppressWarnings(library(rjson))

search_in = fromJSON(search_in_string, simplify = T)
search_out = fromJSON(search_out_string, simplify = T)

# load the experimental design
ExpDesign <- read.table(search_in["exp_design_file"][[1]],sep="\t",header=T)

rm(search_in_string, search_out_string)


## assuming that the computer allows min. 4 threads
NumThreads <- 4

# library causes the execution to fail if the library is missing
suppressWarnings(suppressMessages(library(lattice)))
suppressWarnings(suppressMessages(library(stringr)))
suppressWarnings(suppressMessages(library(mzID)))
suppressWarnings(suppressMessages(library(MSnbase)))

# warnings as stdout
sink(stdout(), type = "message")

message("Verifying availability of files ...")

# change to the input directory
spectradirs <- unlist(search_in["input_directory"])
out_dir <- search_in["workdir"][[1]]
#setwd(out_dir)

## Folder names for different runs (e.g. different replicates, ...)
samples <- unique(as.character(ExpDesign$spec_dir))

# If only one folder, then set correct folder names
if(length(samples) == 1) {
    samples <- "./"
    ExpDesign$spec_dir <- "./"
}
sampledirs <- paste(out_dir,"/",samples,sep="")
all_ident_files <- search_out[[1]]
if (length(sampledirs) != length(spectradirs) | length(sampledirs) != length(all_ident_files)) {
    stop("Unequal number of sample folders")
}
names(sampledirs) <- names(spectradirs) <- names(all_ident_files) <- samples

message("OK")
```

```
Verifying availability of files ...
OK
```

##### Load and Quantify Spectra at the peptide level¶

In [10]:

```
display(HTML("<h4>Processing identification data...</h4>"))

def loadPsms(s):
    """
    Loads the PSMs of the specified samples in the R environment.
    :param s: The sample to be processed
    :return: Nothing is returned. A "psms" object is created in the R environment.
    """
    %R -i s
    
    ro.reval('''
    # warnings as stdout
    sink(stdout(), type = "message")
    
    ident_files <- all_ident_files[s]
    
    if (is.null(ident_files)) stop("Error: No identification files")
    
    if (!file.exists(ident_files)) stop(paste("Error: Cannot find identification files", ident_files))
    
    # ---- load the PSMs ----
    max_fdr <- search_in["target_fdr"]
    psms <- read.csv(ident_files, sep = "\t",stringsAsFactors = F)
    
    if (! "Decoy" %in% names(psms)) {
        stop("Error: No decoy information available in output file")
    }
    
    num_psms <- nrow(psms)
    ''')
    
    num_psms = ro.r.num_psms[0]
    display(HTML(" - Loaded " + str(num_psms) + " PSMs"))
    
    ro.reval('''
    # ---- Confidence filter ----
    # TODO: This could be replaced by the PeptideShaker functionality
    psms <- psms[order(psms[, "Confidence...."], decreasing = T), ]
    decoy.psms <- which(psms[, "Decoy"] == "1")
  
    decoy.count <- 0
  
    for (decoy.index in decoy.psms) {
        decoy.count <- decoy.count + 1
        target.count <- decoy.index - decoy.count
        cur.fdr <- (decoy.count * 2) / (decoy.count + target.count)
    
        if (cur.fdr > max_fdr) {
          # filter
          psms <- psms[1:decoy.index - 1,]
          break
        }
    }
  
    num_psms <- nrow(psms)
    
    if (nrow(psms) < 1) {
        stop("Error: No valid PSMs found")
    }
    ''')
    
    num_psms = ro.r.num_psms[0]
    max_fdr = ro.r.max_fdr[0][0]
    display(HTML(" - Filtered " + str(num_psms) + " PSMs @ " + str(max_fdr) + " FDR"))
    
    ro.reval('''
    # ---- prepare for MSnbase ----
    psms$rank <- 1
    psms$desc <- psms$Protein.s.
    psms$spectrumID <- psms$Spectrum.Title 
    psms$spectrumFile <- psms$Spectrum.File
    psms$idFile <- ident_files
    # remove unnecessary PTMs from modified sequence
    # TODO: define whether to take oxidation, ...
    psms$Modified.Sequence <- gsub("<cmm>","",psms$Modified.Sequence)
    psms$Modified.Sequence <- sub("[a-z,A-Z]*-","",psms$Modified.Sequence)
    psms$Modified.Sequence <- sub("-[a-z,A-Z]*","",psms$Modified.Sequence)
    psms$Modified.Sequence <- gsub("<iTRAQ>","",psms$Modified.Sequence)
    psms$Modified.Sequence <- gsub("<TMT>","",psms$Modified.Sequence)
    psms$sequence <- psms$Modified.Sequence
    ''')
    
def loadSpectra(s):
    """
    Loads the spectra from the MGF files and stores them
    in a list of MSnbase Exp
    :param s: The sample to process
    :return: None, a list called "allSpectra" is created in the R environment holding
             one object per MGF file
    """
    display(HTML(" - Loading spectra (this may take a while) ..."))
    
    ro.reval('''
    # --- Load the spectra ----
    mgf_path <- paste0(search_in[["workdir"]], "/", gsub("/$", "", s)[[1]])
    mgf_files <- list.files(mgf_path,pattern="mgf.filtered$",full.names = T)
    if (is.null(mgf_files) || length(mgf_files) < 1)   stop("Error: No spectrum files")
    
    for (mgf_file in mgf_files) {
        if (!file.exists(mgf_file)) {
             stop("Error: Cannot find mgf file ", mgf_file)       
        }
    }
    
    allSpectra <- list()
    for (mgf_file in mgf_files) {
        invisible(capture.output(myExp1 <- readMgfData(mgf_file, verbose=F)))
    
        for (i in 1:ncol(fData(myExp1))) {
          if (is.factor(fData(myExp1)[,i]))
            fData(myExp1)[,i] <- as.character(fData(myExp1)[,i])
        }
        fData(myExp1)$spectrumFile <- mgf_file
        allSpectra[[mgf_file]] <- myExp1
    }
    ''')

def quantify_mgf_file(mgf_file):
    """
    Quantify the reporter peaks in the given MGF file and merge the
    identification data.
    :param mgf_file: name of the MGF file to quantify
    :return None: Changes
    """
    %R -i mgf_file
    ro.reval('''
    cl <- makeCluster(NumThreads)
    myExp[[mgf_file]] <- addIdentificationData(
        allSpectra[[mgf_file]],
        psms,
        decoy="Decoy",
        rank="rank",
        acc="Protein.s.",
        icol="Spectrum.Title",
        fcol="TITLE",
        desc="desc",
        pepseq="Modified.Sequence",
        verbose=F)

    # quantify everything
    qnt[[mgf_file]] <- quantify(myExp[[mgf_file]], 
                                method = "sum", 
                                reporters = quant.method, 
                                strict = F, verbose = F)
    stopCluster(cl)
    ''')

%R PSMDat <- PepDat <- ProtDat <- list()
for s in samples:
    # Organize technical runs in the same folder
    # filenames
    %R -i s
    display(HTML("<i>Processing files from folder " + str(s) + " (" + str(samples.index(s)+1) + " of " + str(len(samples)) + ")</i>" ))
    
    cache_file_path = os.path.join(search_in["workdir"], s, "RawQuant.rds")
    %R -i cache_file_path
    
    if os.path.isfile(cache_file_path):
        display(HTML("Loading data from cache..."))
        # load the cache data
        %R load(cache_file_path)
        mgf_files = []
        mgf_files.extend(ro.r.mgf_files)
    else:
        loadPsms(s)

        loadSpectra(s)

        # get the quantification method
        ro.reval('''
        known.methods <- c("TMT10","TMT6","iTRAQ4","iTRAQ8","iTRAQ4", "iTRAQ8")
        selected.method <- gsub(" \\\\(.*", "", search_in["labelling_method"][[1]])

        if (!selected.method %in% known.methods) {
            stop("Error: Labelling method not supported")
        }

        quant.method <- get(selected.method)

        # run the merging and quantification in parallel
        myExp <- qnt <- list()
        ''')

        # quantify every MGF file and add the id data
        mgf_files = []
        mgf_files.extend(ro.r.mgf_files)
        num_mgfs = str(len(mgf_files))
        for mgf_file in mgf_files:
            display(HTML("Getting reporter ions (quantification) from " + mgf_file + "\n(" + 
                         str(mgf_files.index(mgf_file)+1) + " of " +  num_mgfs + " files)"))

            quantify_mgf_file(mgf_file)
        
        # save the data to the cache
        %R save(myExp, qnt, mgf_files, quant.method, file = cache_file_path)

    # create the Q/C plots
    for mgf_file in mgf_files:
        %R -i mgf_file
        
        # Show the Q/C plot
        ro.reval('''
        # Plot reporter QC
        plotQCHist <- function() {
        hist(unlist(mz(myExp[[mgf_file]])), 1000, main=paste("m/z accuracy of reporter ions\n",mgf_file), 
            xlim=range(quant.method@mz)+c(-1,1),xlab="m/z",col="#666666",border=NA)
            abline(v=quant.method@mz,col=2,lwd=0.5)
        }
        # save histograms as pdf and png files       
        png(filename=paste(sampledirs[s],"/QC_ReporterIons_",basename(mgf_file),".png",sep=""),width=1200,height=1200)
        plotQCHist()
        dev.off()
        pdf(file=paste(sampledirs[s],"/QC_ReporterIons_",basename(mgf_file),".pdf",sep=""),width=8,height=8)
        plotQCHist()
        dev.off()

        ''')
        %R --width=1000 plotQCHist()
        
    # perform the impurity correction and normalisation
    display(HTML("Normalising PSM features..."))
    ro.reval('''    
    imp <- makeImpuritiesMatrix(length(quant.method),edit=F)
    for (i in 1:length(qnt)) {
        qnt[[i]] <- purityCorrect(qnt[[i]],imp)
        
        n.features.before <- nrow(qnt[[i]])
    
        # remove features missing in 50% of the samples
        qnt[[i]] <- filterNA(qnt[[i]], pNA=0.5)
        qnt[[i]] <- filterZero(qnt[[i]], 0.5)
    
        message("  Removed ", n.features.before - nrow(qnt[[i]]), " features with 50% missing values")
        
        # remove all unidentified spectra
        qnt[[i]] <- removeNoId(qnt[[i]])

        # MSnbase function generates unwanted negative values: qnt[[i]] <- normalise(qnt[[i]],"center.median")
        
        # log-transformation
        tdat <- log2(exprs(qnt[[i]]))

        # set -Inf to NA
        tdat[tdat == -Inf] <- NA

        # normalize by subtracing col-wise median
        exprs(qnt[[i]]) <- t(t(tdat) - apply(tdat, 2, median, na.rm=T))
                
        # update feature names in preparation of merging
        qnt[[i]] <- updateFeatureNames(qnt[[i]],label = paste("Sample",i))
    }
    ''')
    
    # combine the quant data
    display(HTML("Merging quantification data..."))
    ro.reval('''
    # ---- combine the quantification data ----
    names(qnt) <- NULL
    allqnt <- do.call("combine",args=qnt)

    # ---- add the sample annotations ----
    SampleExpDesign <- ExpDesign[ExpDesign$spec_dir == s,]
    if (nrow(SampleExpDesign) != nrow(pData(allqnt))) {
        stop("Error: Experimental design does not fit the number of quantified samples.")
    }

    # merge the annotations
    pdata.org <- pData(allqnt)
    pdata.combined <- merge(pdata.org, SampleExpDesign, all.x = T, all.y = F, by.x = 0, by.y = "channel")
    rownames(pdata.combined) <- pdata.combined[, "Row.names"]
    pdata.combined$Row.names <- NULL
    outfile <- paste(sampledirs[s],"/AllQuantPSMs.RData",sep="")
    ''')
    
    # save the quant data
    display(HTML("\r - Saving PSM quantifications to " + str(ro.r.outfile[0])))
    ro.reval('''
    # save
    pData(allqnt) <- pdata.combined[colnames(allqnt), ]
  
    ## Setting the stage for the iPQF inference method
    names(fData(allqnt))[which(names(fData(allqnt))=="Protein.s.")] <- "accession"
    names(fData(allqnt))[which(names(fData(allqnt))=="Variable.Modifications")] <- "modifications"
    names(fData(allqnt))[which(names(fData(allqnt))=="m.z")] <- "mass_to_charge"
    names(fData(allqnt))[which(names(fData(allqnt))=="Confidence....")] <- "search_engine_score"
    
    
    write.csv(cbind(fData(allqnt),exprs(allqnt)),paste(sampledirs[s],"/AllQuantPSMs.csv",sep=""))  
    save(allqnt, file= outfile)
    
    PSMDat[[s]] <- allqnt 
    ''')
display(HTML("<b>Finished successfully</b>"))
```

###### Processing identification data...

*Processing files from folder ./ (1 of 1)*

Loading data from cache...

Normalising PSM features...

```
  Removed 753 features with 50% missing values

  Removed 248 features with 50% missing values

  Removed 252 features with 50% missing values

  Removed 212 features with 50% missing values

  Removed 731 features with 50% missing values

  Removed 277 features with 50% missing values

  Removed 120 features with 50% missing values

  Removed 772 features with 50% missing values

  Removed 379 features with 50% missing values

  Removed 354 features with 50% missing values

  Removed 444 features with 50% missing values

  Removed 277 features with 50% missing values

  Removed 334 features with 50% missing values
```

Merging quantification data...

- Saving PSM quantifications to /home/biodocker/OUT/.//AllQuantPSMs.RData

**Finished successfully**

##### Basic Quality Control¶

In [11]:

```
display(HTML("<h4>Processing PSMs ...</h4>"))

%R condNames  <- list()

ro.reval('''
# Used panel functions
panel.cor <- function(x, y, digits=2, prefix="", cex.cor, ...) 
{
    usr <- par("usr"); on.exit(par(usr)) 
    par(usr = c(0, 1, 0, 1)) 
    r <- abs(cor(x, y, use="na.or.complete")) 
    txt <- format(c(r, 0.123456789), digits=digits)[1] 
    txt <- paste(prefix, txt, sep="") 
    cex.cor <- 0.8/strwidth(txt) 
    test <- cor.test(x,y) 
    # borrowed from printCoefmat
    Signif <- symnum(test$p.value, corr = FALSE, na = FALSE, 
                  cutpoints = c(0, 0.001, 0.01, 0.05, 0.1, 1),
                  symbols = c("***", "**", "*", ".", " ")) 
 
    text(0.5, 0.5, txt, cex = cex.cor * r) 
}

panel.hist <- function(x, hist.col="#993333", ...)
{
    usr <- par("usr"); on.exit(par(usr))
   par(usr = c(usr[1:2], 0, 1.5) )
    h <- hist(x, plot = FALSE, border=NA,breaks=50)
    breaks <- h$breaks; nB <- length(breaks)
  y <- h$counts; y <- y/max(y)
    rect(breaks[-nB], 0, breaks[-1], y,col=hist.col,border=NA)
}

suppressWarnings(suppressMessages(library(plotly)))
suppressWarnings(suppressMessages(library(reshape)))
''')


for s in samples:
    %R -i s
    display(HTML("<i>Making figures for files from folder " + str(s) + "(" + str(samples.index(s)+1) + " of " + str(len(samples)) + ")</i>" ))
    ro.reval('''
    allqnt <- PSMDat[[s]]
    
    condNames[[s]] <- ExpDesign[ExpDesign$spec_dir == s,"sample_orig"]
    conditions <- condNames[[s]]
    if (sampledirs[s] != "")
    if(s != "./" & length(sampledirs) > 1) condNames[[s]] <- paste(paste("Sample", 1:length(conditions)), condNames[[s]], s, sep="\n")
    pData(allqnt)$sample_name <- paste(condNames[[s]], sampledirs[s])
    pData(allqnt)$sample_group <- condNames[[s]]
    
    
    # Better naming:
    samplename <- ifelse(s=="./","",s)
    
    ## QC Plots
    # violin plots to show distribution of PSM quantifications
    conditionsPlot <- rep(conditions, each=nrow(exprs(allqnt)))
    p <- ggplot(melt(exprs(allqnt)), aes(x=X2, y=value, fill=conditionsPlot)) + geom_violin(trim=FALSE) + 
    geom_boxplot(width=0.1) + theme_classic() + theme(axis.text.x = element_text(angle = 90, hjust = 1)) +
    xlab("") + ylab("PSM expression") + ggtitle(paste("Distributions in MS run",samplename))
    png(filename=paste(sampledirs[s],"/QC_PSM_violinplots.png",sep=""),width=600,height=600)
    print(p)
    dev.off()
    pdf(file=paste(sampledirs[s],"/QC_PSM_violinplots.pdf",sep=""),width=8,height=8)
    print(p)
    dev.off()
    
    ## histograms of PSMs and peptides per peptide/protein, counting for unique and shared
    QC_PSMHist <- function() {
            par(mfrow=c(2,3))
# all PSMs per peptide
hist(table(fData(allqnt)$Sequence),xlab="PSMs per peptide",100,border=0,col="#AA4444", main="PSMs per peptide")

    # all PSMs per protein (unique and shared)
AllSingleProts <- gsub(" ","",unlist(strsplit(fData(allqnt)$accession,",")))
hist(table((AllSingleProts)),xlab="PSMs per protein",100,border=0,col="#44AA44", main="Shared and unique PSMs per protein")

# all PSMS per protein group
hist(table(fData(allqnt)$accession),xlab="PSMs per protein group",100,border=0,col="#4444AA", main="Unique PSMs per protein group")

PeptideCounter <- unique(fData(allqnt)[,c("Sequence","accession")])
# unique and shared peptides per protein
AllSingleProts <- gsub(" ","",unlist(strsplit(PeptideCounter$accession,",")))
hist(table((AllSingleProts)),xlab="Peptides per protein",100,border=0,col="#AAAA44", main="Shared and unique peptides per protein")

# all unique peptides per protein group
hist(table(PeptideCounter$accession),xlab="Peptides per protein group",100,border=0,col="#AA44AA", main="Unique peptides per protein group")

# proteins per peptide
hist(sapply(strsplit(PeptideCounter$accession,","), length),xlab="Proteins matching peptide",100,border=0,col="#44AAAA", main="Proteins in protein groups")
    }
    png(filename=paste(sampledirs[s],"/QC_PSM_and_peptide_distribution.png",sep=""),width=800,height=500)
    QC_PSMHist()
    dev.off()
    pdf(file=paste(sampledirs[s],"/QC_PSM_and_peptide_distribution.pdf",sep=""),width=14,height=8)
    QC_PSMHist()
    dev.off()   
     
    par(mfrow=c(1,1))
    # Pairwise comparison of samples within on MS run
    QC_Pairs <- function () { 
      pairs(exprs(allqnt),lower.panel=panel.smooth, upper.panel=panel.cor, diag.panel=panel.hist, 
        main = paste("MS run",samplename),
        cex=0.1,col="#33333388",pch=15, labels =  condNames[[s]])
    }
    png(filename=paste(sampledirs[s],"/QC_Pairwise_comparison.png",sep=""),width=800,height=800)
    QC_Pairs()
    dev.off()
    pdf(file=paste(sampledirs[s],"/QC_Pairwise_comparison.pdf",sep=""),width=15,height=15)
    QC_Pairs()
    dev.off()   

    ''')
    # display figures
    
    %R --width 800  print(p);QC_PSMHist();par(mfrow=c(1,1));QC_Pairs()

    ro.reval('''
    PSMDat[[s]] <- allqnt    
    ''')
```

###### Processing PSMs ...

*Making figures for files from folder ./(1 of 1)*

##### Protein Inference¶

In [12]:

```
display(HTML("<h4>Protein quantification ...</h4>"))

ro.reval('''
summarization_function <- search_in["summarization_method"][[1]]
    
# get minimum peptides
min_peps <- search_in["min_protein_psms"][[1]]        
''')

display(HTML("Using " + str(ro.r.summarization_function[0]) + " for protein summarization"))
display(HTML("and requiring a minimum of " + str(ro.r.min_peps[0]) + " PSMs per protein"))

for s in samples:
    %R -i s
    display(HTML("<i>Making figures for files from folder " + str(s) + " (" + str(samples.index(s)+1) + " of " + str(len(samples)) + ")</i>" ))
    display(HTML("Filtering and summarization"))
    ro.reval('''
    load(paste(sampledirs[s],"/AllQuantPSMs.RData",sep=""))
    
    allqnt <- PSMDat[[s]]
    
    ### TODO: remove only previously specified variable PTMs
    # Remove other PTMs for quantification
    if (!search_in$use_ptms_for_quant) {
        message(paste0("Removing ",sum(fData(allqnt)$modifications !="") ," modified sequences"))
        allqnt <- allqnt[fData(allqnt)$modifications == "",]
    }

    allProts <- NULL
    # remove missing values for iPQF
    if(summarization_function == "iPQF") {
        exprs(allqnt) <- 2^exprs(allqnt)
        exprs(allqnt)[exprs(allqnt) == -Inf] <- NA
        
        # remove all spectra with missing values
        has.missing.values <- (rowSums(is.na(exprs(allqnt))) > 0 | rowSums(exprs(allqnt) == 0)) > 0    
        message(paste0("Removing ", sum(has.missing.values), " PSMs with missing values for quantification"))
        allqnt <- allqnt[!has.missing.values, ]
           
        # filter for minimal peptide number
        has_too_few_peps <- fData(allqnt)$npep.prot < min_peps
        num_prot_rm <- length(unique(fData(allqnt)[has_too_few_peps, "accession"]))
        
        allqnt <- allqnt[!has_too_few_peps,]        
        
        message(paste("Removing",sum(has_too_few_peps),"PSMs corresponding to",num_prot_rm,"proteins with less then",min_peps,"peptides"))

        allProts <- combineFeatures(allqnt, fun=summarization_function,
                                groupBy=fData(allqnt)$accession, 
                                ratio.calc = "none", method.combine = T,
                                verbose=F)
        exprs(allProts) <- log2(exprs(allProts))
    } else {
    
        # remove all spectra with missing values
        has.missing.values <- rowSums(is.na(exprs(allqnt)) | rowSums(exprs(allqnt) == 0, na.rm=T)) > ncol(allqnt)*0.5    
        message(paste0("Removing ", sum(has.missing.values), " PSMs with more than 50% missing values for quantification"))
        allqnt <- allqnt[!has.missing.values, ]

        # filter for minimal peptide number
        has_too_few_peps <- fData(allqnt)$npep.prot < min_peps
        num_prot_rm <- length(unique(fData(allqnt)[has_too_few_peps, "accession"]))
        
        allqnt <- allqnt[!has_too_few_peps,]
        
        message(paste("Removing",sum(has_too_few_peps),"PSMs corresponding to",num_prot_rm,"proteins with less then",min_peps,"peptides"))

        allProts <- combineFeatures(allqnt, na.rm=T, fun=summarization_function,
                                groupBy=fData(allqnt)$accession, 
                                verbose=F)
    }
    ''')
    display(HTML("Creating figure(s)"))

    ro.reval('''
    # create a plot of all protein expression values
    p <- ggplot(melt(exprs(allProts)), aes(x=X2, y=value, fill=rep(conditions,each=nrow(exprs(allProts))))) + 
            geom_violin(trim=FALSE) + 
            geom_boxplot(width=0.1) +
            theme_classic() + 
            theme(axis.text.x = element_text(angle = 90, hjust = 1)) +
            xlab("") + 
            ylab("Protein expression") + 
            guides(fill=guide_legend(title="Conditions"))

    # save the plot
    png(filename=paste(sampledirs[s],"/QC_Protein_violinplots.png",sep=""),width=800,height=800)
    print(p)
    dev.off()
    pdf(file=paste(sampledirs[s],"/QC_Protein_violinplots.pdf",sep=""),width=8,height=8)
    print(p)
    dev.off()   
    ''')
    
    # Display plot
    %R --width 800 print(p)

    ro.reval('''
    # save the expression values
    write.exprs(allProts,file=paste(sampledirs[s],"/AllQuantProteins.csv",sep=""))
  
    ProtDat[[s]] <- allProts <- exprs(allProts)
    colnames(ProtDat[[s]]) <- condNames[[s]]
    ''')
```

###### Protein quantification ...

Using medpolish for protein summarization

and requiring a minimum of 2.0 PSMs per protein

*Making figures for files from folder ./ (1 of 1)*

Filtering and summarization

```
Removing 11717 modified sequences

Removing 40 PSMs with more than 50% missing values for quantification

Removing 17892 PSMs corresponding to 2314 proteins with less then 2 peptides

Your data contains missing values. Please read the relevant section in
the combineFeatures manual page for details the effects of missing
values on data aggregation.
```

Creating figure(s)

##### Sample similarity and statistics¶

In [13]:

```
display(HTML("<h4>Protein quantification ...</h4>"))

display(HTML("Merging sample runs from different folders ..."))
ro.reval('''
par(mfrow=c(1,1))

# Adjust each protein of each MS run by the mean of their expressions over the channels
# or reference condition (e.g. pool)?
    
for (s in samples) {
    norm_vec <- 0
    if (search_in["ref_condition"] == "Mean over channels") {
        norm_vec <- rowMeans(ProtDat[[s]],na.rm=T)
    } else {
        ref_cols <- which(ExpDesign$sample_group[ExpDesign$spec_dir == s] == search_in["ref_condition"])
        if (length(ref_cols) > 0) {
            norm_vec <- rowMeans(ProtDat[[s]][, ref_cols, drop=F],na.rm=T)
        } else {
            message(paste("No reference samples in MS run",s,"! Taking mean of all channels to adjust relative protein abundances."))
            norm_vec <- rowMeans(ProtDat[[s]],na.rm=T)        
        }
    }
    ProtDat[[s]] <- ProtDat[[s]] - norm_vec
}


# Merge different samples
    totProts <- ProtDat[[1]]
if (length(samples)>1) {
    print("Merging samples (if needed) ...")
    totProts <- data.frame(rownames(totProts),totProts)
    for (s in samples[2:length(samples)])
        totProts <- merge(totProts,ProtDat[[s]], all=T, by.x=1, by.y=0)
    rownames(totProts) <- totProts[,1]
    totProts <- totProts[,2:ncol(totProts)]
} 
write.csv(totProts,file=paste(out_dir,"/AllQuantProteinsInAllSamples.csv",sep=""))
''')

display(HTML("Principal component analysis ..."))
ro.reval('''
# PCA
pca <- princomp((totProts[complete.cases((totProts)),]))
  #plot(pca)

QC_PCA <- function() {
    plot(pca$loadings, cex=2, pch=16, col=as.numeric(as.factor(ExpDesign[["sample_orig"]])))
    text(pca$loadings,colnames(totProts), pos=2)
}
png(filename=paste(out_dir,"/QC_Stat_PCA.png",sep=""),width=800,height=800)
QC_PCA()
dev.off()
pdf(file=paste(out_dir,"/QC_Stat_PCA.pdf",sep=""),width=8,height=8)
QC_PCA()
dev.off()
num_prots <- nrow(allProts)
''')
# display figure
%R --width 1000 QC_PCA()

display(HTML("Quantified a total of " + str(ro.r.num_prots[0]) + " protein groups"))

display(HTML("Running LIMMA for statistical tests ..."))

ro.reval('''
##Statistics
library(limma)
library(qvalue)
NumCond <- length(unique(ExpDesign$sample_orig))
  if (NumCond < 2)
      stop("Only 1 experimental condition -> no statistics")

design <- model.matrix(~0+factor(ExpDesign$sample_group)-1)
  colnames(design)<-make.names(paste(unique(ExpDesign$sample_orig),sep=""))
  contrasts<-NULL
  First <- which(unique(ExpDesign$sample_group) == search_in[["stat_condition"]])
  for (i in (1:NumCond)[-First]) contrasts<-append(contrasts,paste(colnames(design)[i],"-",colnames(design)[First],sep=""))
  print(paste("Statistical tests carried out to compare:",contrasts))
  contrast.matrix<-makeContrasts(contrasts=contrasts,levels=design)
  # print(dim(Data))
  lm.fitted <- lmFit(totProts,design)
  lm.contr <- contrasts.fit(lm.fitted,contrast.matrix)
  lm.bayes <- eBayes(lm.contr)
  #topTable(lm.bayes)
 plvalues <- lm.bayes$p.value
fcs <- lm.bayes$coefficients
  qlvalues <- matrix(NA,nrow=nrow(plvalues),ncol=ncol(plvalues),dimnames=dimnames(plvalues))
  # qvalue correction
 for (i in 1:ncol(plvalues)) {
    tqs <- qvalue(na.omit(plvalues[,i]))$qvalues
    qlvalues[names(tqs),i] <- tqs
  }
  

par(mfrow=c(1,3))

statTable <- NULL

# Visualizations: volcano plot, number of regulated proteins per FDR, interactive table

# p-value distributions

QC_pvals <- function(i) {
    hist(plvalues[,i],100,border=NA,col="#555555", main=paste("p-value distribution\nComparison",colnames(fcs)[i], sep="\n"),         
        xlab="p-values")
}
# volcano plots
QC_volcanos <- function(i) {
    plot(fcs[,i], -log10(qlvalues[,i]),pch=16,col="#44FFFF99",
        xlab="log(fold-change)", ylab="-log10(FDR)", main=paste("Volcano plot\nComparison",
                                                            colnames(fcs)[i], sep="\n"),
        ylim=c(0,max(2,max(-log10(qlvalues[,i]), na.rm=T))))
    abline(h=-log10(c(0.001,0.01,0.05)), col=2:4,lwd=2)
    text(c(0,0,0), c(3,2,1.3),c("FDR=0.001","FDR=0.01","FDR=0.05"),pos=1)
}
# Number of sigificant features vs. FDR
QC_NumSig <- function(i) {
    ddd <- c(0.0001,sort(qlvalues[,i]))
    plot(ddd, 0:(length(ddd)-1),type="l",xlim=c(1e-3,1),log="x", main=paste("Significant protein versus FDR\nComparison",colnames(fcs)[i], sep="\n")) 
    abline(v=c(0.001,0.01,0.05), col=2:4, xlab="FDR", ylab="Number of proteins")
}
QC_AllStats <- function(i) {
    QC_pvals(i)
    QC_volcanos(i)
    QC_NumSig(i)
}

for (i in 1:ncol(plvalues)) {    
    # export figures to pdf/png
    png(filename=paste(out_dir,"/QC_Stat_Summary_",colnames(fcs)[i],".png",sep=""),width=1000,height=500)
    par(mfrow=c(1,3))
    QC_pvals(i)
    QC_volcanos(i)
    QC_NumSig(i)   
    dev.off()
    pdf(file=paste(out_dir,"/QC_Stat_Summary_",colnames(fcs)[i],".pdf",sep=""),width=15,height=8)
    par(mfrow=c(1,3))
    QC_pvals(i)
    QC_volcanos(i)
    QC_NumSig(i)   
    dev.off()
par(mfrow=c(1,1))    
    
}
''')

display(HTML("Saving statistics results ..."))
ro.reval('''
# How far should we go? Clustering (when having more then 2 groups)? 
# Hierarchical clustering of significant features? clusterProfiler?


# Saving stats
statOut <- cbind(fcs, plvalues, qlvalues)
colnames(statOut) <- paste(rep(c("log(fold-change)","p-values","q-values"), each=ncol(plvalues)), colnames(statOut))
statOut <- cbind(Proteins=rownames(fcs),statOut)
#print(head(statOut))
write.csv(statOut, paste(out_dir,"/DifferentiallyRegulatedProteins.csv",sep=""),row.names=F)

''')

%R -w 1000  aa<-c(1,3); bb <- ncol(plvalues); par(mfrow=aa); for (i in 1:bb) QC_AllStats(i)
```

###### Protein quantification ...

Merging sample runs from different folders ...

Principal component analysis ...

Quantified a total of 1432 protein groups

Running LIMMA for statistical tests ...

```
Attaching package: 'limma'


The following object is masked from 'package:BiocGenerics':

    plotMA


[1]
 "Statistical tests carried out to compare: Treatment.1-Control"
```

Saving statistics results ...

##### Run your own script¶

In [14]:

```
%%R --res 150 --width 700 --height 1000
# spiked proteins
spiked.proteins <- c("ENO1" = "P06733", "ARG1" = "P05089", "FABP4" = "P15090", "PTGES3" = "Q15185", "KPNA2" = "P52292", 
                     "LASP1" = "Q14847", "HDAC3"= "O15379", "KCNIP3" = "Q9Y2W7", "OTUB1" = "Q96FW1", "GABARAPL1" = "Q9H0R8",
                     "GAS7" = "O60861", "EZR" = "P15311", "Alb" = "P02768")

# Background fold-changes

lfc <- lm.bayes$coefficients
boxplot(lfc[!rownames(lfc) %in% as.character(spiked.proteins),], 
        ylab = "log-fold change",
       main = "Fold-change of background proteins")
```

In [15]:

```
%%R --width 800 -r 100 --height 800

proteins <- read.csv("OUT/AllQuantProteins.csv", sep = "\t", row.names = 1)

# proteins <- read.csv("OUT/AllQuantProteinsInAllSamples.csv", row.names = 1)

proteins <- proteins[rownames(proteins) %in% as.character(spiked.proteins), ]

# change to gene names
name.index <- names(spiked.proteins)
names(name.index) <- as.character(spiked.proteins)
rownames(proteins) <- name.index[rownames(proteins)]

# load the spiked in measurements
spiked.conc <- read.csv("data/spiked_proteins.tsv.csv", sep = "\t", dec = ",", row.names = 1)

# compare all proteins
plot.data <- data.frame()
for (gene in names(spiked.proteins)) {
    plot.data <- rbind(plot.data, data.frame(
        gene = gene,
        spiked = log2(as.numeric(spiked.conc[gene, ])),
        measured = as.numeric(proteins[gene, ])
    ))
}

library(ggplot2)
ggplot(plot.data, aes(x = spiked, y = measured, group = gene, color = gene)) +
    geom_point() +
    geom_line() +
    facet_wrap(~ gene) +
    labs(x = "Spiked in concentration", y = "Measured protein abundance", color = "Protein")
```

In [16]:

```
%%R --width 600 --height 300 --res 100

getFC <- function(protein, cols.grp1, cols.grp2) {
    grp1 <- as.numeric(proteins[protein, cols.grp1])
    grp2 <- as.numeric(proteins[protein, cols.grp2])
    
    #print(paste0("Group 1: ", paste(grp1, collapse = ", ")))
    #print(paste0("Group 2: ", paste(grp2, collapse = ", ")))
    
    mean.grp1 <- mean(2^grp1)
    mean.grp2 <- mean(2^grp2)
    
    #print(paste0("Mean group 1: ", mean.grp1))
    #print(paste0("Mean group 2: ", mean.grp2))
    
    #return(abs(log2(mean.grp1 / mean.grp2)))
    return(mean.grp2 / mean.grp1)
}

generateFcForGene <- function(gene, cols1, cols2, cols4, col20, col40) {
    logfc.values <- data.frame()
    logfc.values <- rbind(logfc.values, data.frame(gene = gene, real.fc = 2, our.fc = getFC(gene, cols1, cols2)))
    logfc.values <- rbind(logfc.values, data.frame(gene = gene, real.fc = 4, our.fc = getFC(gene, cols1, cols4)))
    logfc.values <- rbind(logfc.values, data.frame(gene = gene, real.fc = 20, our.fc = getFC(gene, cols1, col20)))
    logfc.values <- rbind(logfc.values, data.frame(gene = gene, real.fc = 40, our.fc = getFC(gene, cols1, col40)))
    
    #logfc.values <- rbind(logfc.values, data.frame(gene = gene, real.fc = 2, our.fc = getFC(gene, cols2, cols4)))
    #logfc.values <- rbind(logfc.values, data.frame(gene = gene, real.fc = 10, our.fc = getFC(gene, cols2, col20)))
    #logfc.values <- rbind(logfc.values, data.frame(gene = gene, real.fc = 20, our.fc = getFC(gene, cols2, col40)))
    #
    #logfc.values <- rbind(logfc.values, data.frame(gene = gene, real.fc = 5, our.fc = getFC(gene, cols4, col20)))
    #logfc.values <- rbind(logfc.values, data.frame(gene = gene, real.fc = 10, our.fc = getFC(gene, cols4, col40)))
    
    #logfc.values <- rbind(logfc.values, data.frame(gene = gene, real.fc = 2, our.fc = getFC(gene, col20, col40)))
    
    return(logfc.values)
}

#print(proteins["ENO1",])
logfc.values <- data.frame()
logfc.values <- rbind(logfc.values, generateFcForGene("ENO1", c(6,9), c(3,5), c(7, 10), 2, 8))
logfc.values <- rbind(logfc.values, generateFcForGene("ARG1", c(2,4), c(1, 10), c(5, 9), 7, 3))
logfc.values <- rbind(logfc.values, generateFcForGene("FABP4", c(2,3), c(5,8), c(1, 10), 4, 9))
logfc.values <- rbind(logfc.values, generateFcForGene("PTGES3", c(6,9), c(3,4), c(5,8), 1, 10))
logfc.values <- rbind(logfc.values, generateFcForGene("KPNA2", c(7,9), c(4, 10), c(1,8), 5, 2))
logfc.values <- rbind(logfc.values, generateFcForGene("LASP1", c(2, 9), c(1, 10), c(5, 8), 3, 6))
logfc.values <- rbind(logfc.values, generateFcForGene("HDAC3", c(4,6), c(1,2), c(8, 10), 7, 5))
logfc.values <- rbind(logfc.values, generateFcForGene("KCNIP3", c(8,9), c(3,6), c(1, 5), 10, 4))
logfc.values <- rbind(logfc.values, generateFcForGene("OTUB1", c(7, 10), c(1, 6), c(5, 8), 2, 9))
logfc.values <- rbind(logfc.values, generateFcForGene("GABARAPL1", c(3,9), c(1,8), c(5, 10), 4, 7))
logfc.values <- rbind(logfc.values, generateFcForGene("GAS7", c(2,4), c(7,8), c(1,10), 3, 6))
logfc.values <- rbind(logfc.values, generateFcForGene("EZR", c(2,9), c(5,7), c(3,6), 8, 1))

ggplot(logfc.values, aes(x = real.fc, y = our.fc, color = gene, group = gene)) +
geom_point() +
geom_line(data = data.frame(real.fc = c(1, 40), our.fc = c(1, 40)), aes(x = real.fc, y = our.fc), linetype = "dotted", color = "grey30") +
# scale_color_brewer(palette = "Set1") +
labs(color = "Gene", x = "Real fold change", y = "Estimated fold change")

logfc.values$factor.real.fc <- factor(logfc.values$real.fc)

plot.obj <- ggplot(logfc.values, aes(x = factor.real.fc, y = our.fc)) +
geom_boxplot() +
geom_hline(aes(yintercept = real.fc, col = "red")) + 
facet_wrap(~ factor.real.fc, scales = "free", ncol = 4) +
labs(y = "Estimated fold changed", x = "True fold change")
print(plot.obj)

# calculate the RMSE
logfc.values$error <- logfc.values$our.fc - logfc.values$real.fc
logfc.values$error_sq <- logfc.values$error ^ 2

# plot the error distributions
boxplot(logfc.values$error ~ logfc.values$real.fc, xlab = "Real fold-change", ylab = "Error")


result <- do.call("rbind", by(logfc.values, logfc.values$real.fc, function(x) {
    data.frame(
        rmse = sqrt(sum(x$error_sq) / nrow(x)),
        bias = sum(x$error) / nrow(x)
    )
}))

result$rmse_paper <- c(1.0, 1.9, 11.2, 22.3)
result$bias_paper <- c(-0.6, -1.8, -11.7, -21.9)

round(t(result[c("bias", "bias_paper", "rmse", "rmse_paper")]), 1)
#rmse <- by(logfc.values$error_sq, logfc.values$real.fc, function(x) sqrt(sum(x) / length(x)))
#print(rmse )
           
# bias (???)
#bias <- by(logfc.values$error, logfc.values$real.fc, function(x) sum(x) / length(x))
#print(bias)
```

```
              2    4    20    40
bias       -0.6 -2.0 -11.4 -22.7
bias_paper -0.6 -1.8 -11.7 -21.9
rmse        0.6  2.1  11.8  23.5
rmse_paper  1.0  1.9  11.2  22.3
```
