## Supplementary File 2 for "IsoProt: A complete and reproducible workflow to analyse iTRAQ/TMT experiments"

### 2 Workflow

```
In [1]: from button_execute import ExecuteButton
        # when clicking the button, the next 3 Jupyter notebook cells will be executed
        ExecuteButton(button_text="Start Wokflow", n_next_cells=4)
```

```
ExecuteButton(button_text='Start Wokflow', n_next_cells=4, style=ButtonStyle())
```

```
In [2]: %%javascript
        // disable scrolling for all cells
        requirejs("notebook/js/outputarea").OutputArea.prototype._should_scroll = function(lines
```

```
<IPython.core.display.Javascript object>
```

```
In [3]: #%load_ext rpy2.ipython
        get_ipython().run_line_magic('load_ext', 'rpy2.ipython')
```

##### 2.0.1 Specify parameters for database search and evaluation of identified peptide-spectrum matches:

```
In [4]: # This cell contains the complete code to
        # 1.) Display the GUI for the user to enter the search parameters
        # 2.) Launch the search based on these parameters
        # 3.) Display a button once the search is complete to execute the subsequent R analysis

        from shutil import copyfile
        from IPython.display import display

        search_in = None

        def complete_function():
            global search_in

            search_in = search_ui_out.copy()

            # remove the search_ui object
            if "search_ui" in search_in:
                del search_in["search_ui"]

            search_in["on_search_complete"] = search_complete
            %run "Scripts/search.ipy"

        def search_complete():
            # Enable the R analysis button
            btn_analysis.disabled = False
            btn_analysis.button_style = "success"
            print("Start the analysis by clicking the >>Run R Analysis<< button")
```

```
def update_search_parameters(arg=None):
    """
    Update all settings parameters
    """
    global search_ui_out, search_in
    search_ui = search_ui_out["search_ui"]

    if not search_ui:
        raise Exception("Failed to retrieve searchUI object")

    # update the current settings
    search_ui_settings = search_ui.get_settings_as_dict()

    for setting, value in search_ui_settings.items():
        search_in[setting] = value

    # save the settings again
    search_ui.save_config("/home/biodocker/OUT/protocol_parameters.json")

def on_change_input_dir(search_ui, spectra_dir):
    # test if it is our test directory
    test_file = os.path.join(spectra_dir, "iTRAQCancer.mgf")
    exp_design_file = os.path.join(spectra_dir, "exp_design_example.tsv")
    target_exp_design_file = os.path.join("OUT", "exp_design.tsv")

    if "biodocker/IN" in spectra_dir and os.path.isfile(test_file) and os.path.isfile(ex
        # copy the experimental design file to the default location
        copyfile(exp_design_file, target_exp_design_file)
        search_ui.load_exp_design()

# ----------- ENTRY POINT ------------
search_ui_in = {"on_complete_description": "Run my search", "on_complete_function": comp
            "on_change_input_dir": on_change_input_dir}

%run "Scripts/search_ui.ipy"

# create the R analysis button
btn_analysis = ExecuteButton(button_text="Run R analysis", n_next_cells=13, disabled = T
btn_analysis.on_click(update_search_parameters)
display(btn_analysis)

# Test if results are already present and show the R analysis button in this case
if os.path.isdir("OUT") and os.access("OUT", os.R_OK) and \
   os.path.isfile("OUT/protocol_parameters.json") and \
   os.path.isfile("OUT/exp_design.tsv"):
    # prepare the objects
    search_in = search_ui_out.copy()
```

```python
        if "search_ui" in search_in:
            del(search_in["search_ui"])

        # Test whether only one directory or multiple directories exist
        if os.path.isfile("OUT/experiment1_test_1_Extended_PSM_Report.txt"):
            # replicate all global variables as if the search was run
            print("Previous results found in OUT.")
            search_out = {"result_files": ["OUT/experiment1_test_1_Extended_PSM_Report.txt"]
            search_complete()
        else:
            result_files = list()

            # Test every directory whether it contains result files
            for file in os.listdir("OUT"):
                file_path = os.path.join("OUT", file)
                if os.path.isdir(file_path) and os.access(file_path, os.R_OK):
                    result_file_path = os.path.join(file_path, "experiment1_test_1_Extended_
                    if os.path.isfile(result_file_path):
                        result_files.append(result_file_path)

            if len(result_files) > 0:
                print(str(len(result_files)) + " result files found.")
                search_out = {"result_files": result_files}
                search_complete()
```

```
Tab(children=(VBox(children=(HTML(value='<h4>Folders and files</h4>'), HTML(value='Folder for sp
```

```
ExecuteButton(button_text='Run R analysis', disabled=True, n_next_cells=13, style=ButtonStyle())
```

```
<IPython.core.display.Javascript object>
```

```
<IPython.core.display.Javascript object>
```

```
<IPython.core.display.Javascript object>
```

```
Saving parameters to /home/biodocker/OUT/protocol_parameters.json
Deleting existing files in /home/biodocker/OUT/Sample1
Adapting MGF titles...
['/home/biodocker/data/Sample1/Natalia_TMT6_1_240m_1pt5.mgf']
Extracting reporter peaks...
Creating decoy database...
Start: Creating decoy database...
    Completed in 2 sec
```

```
Start: Creating search parameter file
    Completed in 0 sec
Start: SearchCLI
    Completed in 20:57 min
Start: PeptideShakerCLI processing
    Completed in 1:47 min
Start: ReportCLI (conversion to .tsv)
    Completed in 1:33 min
Search Done.
Deleting existing files in /home/biodocker/OUT/Sample2
Adapting MGF titles...
['/home/biodocker/data/Sample2/Natalia_TMT6_2_240min_1pt5.mgf']
Extracting reporter peaks...
Creating decoy database...
Start: Creating decoy database...
    Completed in 2 sec
Start: Creating search parameter file
    Completed in 0 sec
Start: SearchCLI
    Completed in 27:10 min
Start: PeptideShakerCLI processing
    Completed in 1:37 min
Start: ReportCLI (conversion to .tsv)
    Completed in 1:13 min
Search Done.
Start the analysis by clicking the >>Run R Analysis<< button
Saving parameters to /home/biodocker/OUT/protocol_parameters.json
Saving parameters to /home/biodocker/OUT/protocol_parameters.json
Saving parameters to /home/biodocker/OUT/protocol_parameters.json
Saving parameters to /home/biodocker/OUT/protocol_parameters.json
Saving parameters to /home/biodocker/OUT/protocol_parameters.json
Saving parameters to /home/biodocker/OUT/protocol_parameters.json
Saving parameters to /home/biodocker/OUT/protocol_parameters.json
```

### 2.1 Methods Sections

```
In [47]: %run Scripts/methods_generator.ipy
```

```
HTML(value="<b>Citations are added as PubMed ids</b><br />The mass spectrometry data was analyse
```

#### 2.1.1 Convert Search Settings into R Objects

These cells should not produce any output.

```
In [48]: # Since py2ri cannot convert dict objects, simply save everything R needs as a JSON str
```

```
# remove the callback function first
search_in.pop("on_search_complete", None)

import json
search_in_string = json.dumps(search_in)
search_out_string = json.dumps(search_out)

import rpy2.robjects as ro
from IPython.display import display,HTML
import sys
```

In [49]:
```
%%R -i search_in_string,search_out_string -o samples
# Convert the JSON objects back into "natural" R objects
suppressWarnings(library(rjson))

search_in = fromJSON(search_in_string, simplify = T)
search_out = fromJSON(search_out_string, simplify = T)

# load the experimental design
ExpDesign <- read.table(search_in["exp_design_file"][[1]],sep="\t",header=T)

rm(search_in_string, search_out_string)


## assuming that the computer allows min. 4 threads
NumThreads <- 4

# library causes the execution to fail if the library is missing
suppressWarnings(suppressMessages(library(lattice)))
suppressWarnings(suppressMessages(library(stringr)))
suppressWarnings(suppressMessages(library(mzID)))
suppressWarnings(suppressMessages(library(MSnbase)))

# warnings as stdout
sink(stdout(), type = "message")

message("Verifying availability of files ...")

# change to the input directory
spectradirs <- unlist(search_in["input_directory"])
out_dir <- search_in["workdir"][[1]]
#setwd(out_dir)

## Folder names for different runs (e.g. different replicates, ...)
samples <- unique(as.character(ExpDesign$spec_dir))

# If only one folder, then set correct folder names
if(length(samples) == 1) {
```

```
        samples <- "./"
        ExpDesign$spec_dir <- "./"
    }
    sampledirs <- paste(out_dir,"/",samples,sep="")
    all_ident_files <- search_out[[1]]
    if (length(sampledirs) != length(spectradirs) | length(sampledirs) != length(all_ident_
        stop("Unequal number of sample folders")
    }
    names(sampledirs) <- names(spectradirs) <- names(all_ident_files) <- samples

    message("OK")
```

```
Verifying availability of files ...
OK
```

#### 2.1.2 Load and Quantify Spectra at the peptide level

```
In [50]: display(HTML("<h4>Processing identification data...</h4>"))

         def loadPsms(s):
             """
             Loads the PSMs of the specified samples in the R environment.
             :param s: The sample to be processed
             :return: Nothing is returned. A "psms" object is created in the R environment.
             """
             %R -i s

             ro.reval('''
             # warnings as stdout
             sink(stdout(), type = "message")

             ident_files <- all_ident_files[s]

             if (is.null(ident_files)) stop("Error: No identification files")

             if (!file.exists(ident_files)) stop(paste("Error: Cannot find identification files"

             # ---- load the PSMs ----
             max_fdr <- search_in["target_fdr"]
             psms <- read.csv(ident_files, sep = "\t",stringsAsFactors = F)

             if (! "Decoy" %in% names(psms)) {
                 stop("Error: No decoy information available in output file")
             }

             num_psms <- nrow(psms)
```

```
<IPython.core.display.HTML object>

<IPython.core.display.HTML object>

<IPython.core.display.HTML object>
```

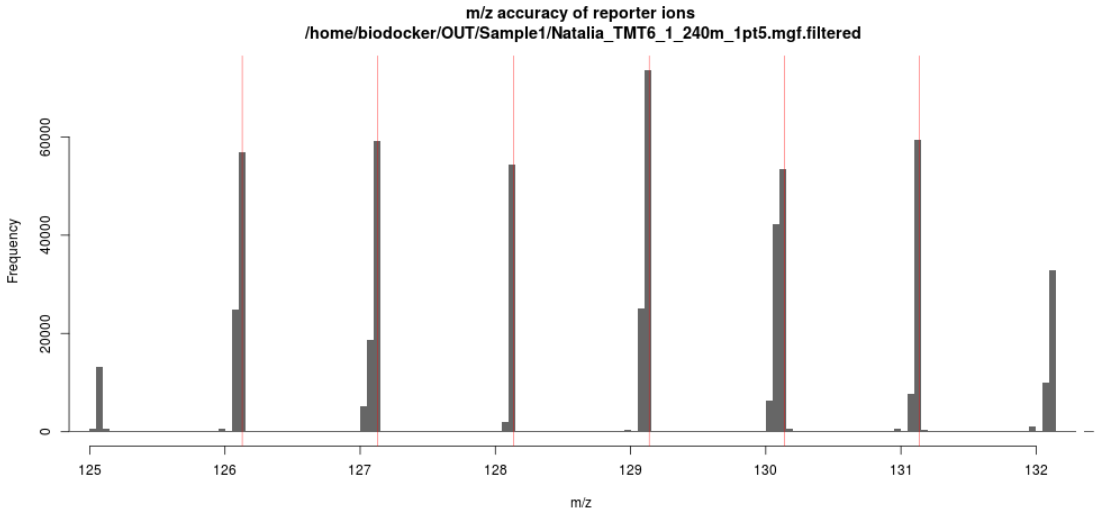


```
<IPython.core.display.HTML object>
```

```
  Removed 160 features with 50% missing values
```

```
<IPython.core.display.HTML object>
```

```
<IPython.core.display.HTML object>
```

```
<IPython.core.display.HTML object>
```

```
<IPython.core.display.HTML object>
```

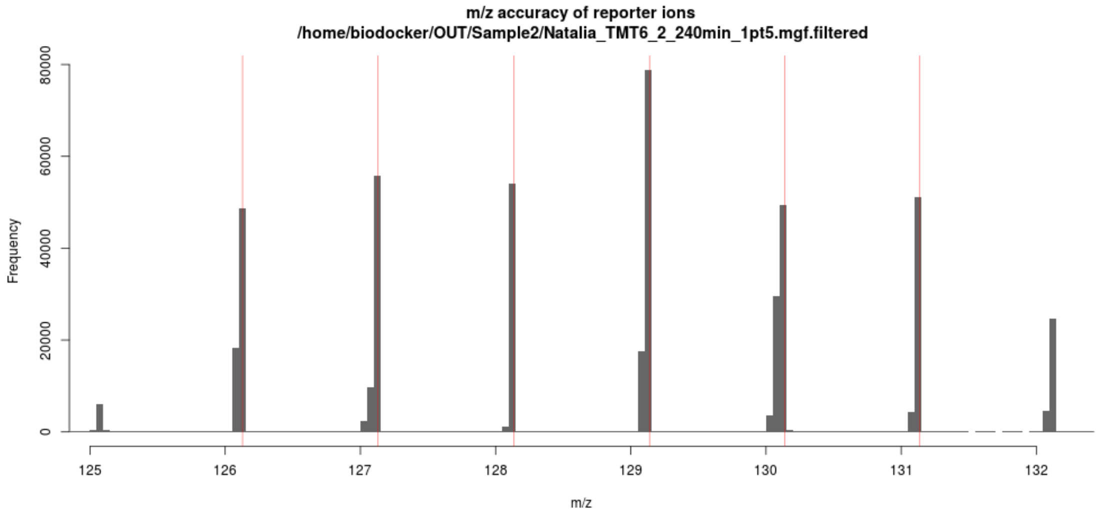


<IPython.core.display.HTML object>

Removed 402 features with 50% missing values

<IPython.core.display.HTML object>

<IPython.core.display.HTML object>

<IPython.core.display.HTML object>

##### 2.1.3 Basic Quality Control

```
In [51]: display(HTML("<h4>Processing PSMs ...</h4>"))

         %R condNames  <- list()

         ro.reval('''
         # Used panel functions
         panel.cor <- function(x, y, digits=2, prefix="", cex.cor, ...)
         {
             usr <- par("usr"); on.exit(par(usr))
             par(usr = c(0, 1, 0, 1))
             r <- abs(cor(x, y, use="na.or.complete"))
             txt <- format(c(r, 0.123456789), digits=digits)[1]
```

```
    txt <- paste(prefix, txt, sep="")
    cex.cor <- 0.8/strwidth(txt)
    test <- cor.test(x,y)
    # borrowed from printCoefmat
    Signif <- symnum(test$p.value, corr = FALSE, na = FALSE,
                  cutpoints = c(0, 0.001, 0.01, 0.05, 0.1, 1),
                  symbols = c("***", "**", "*", ".", " "))

    text(0.5, 0.5, txt, cex = cex.cor * r)
}

panel.hist <- function(x, hist.col="#993333", ...)
{
    usr <- par("usr"); on.exit(par(usr))
   par(usr = c(usr[1:2], 0, 1.5) )
    h <- hist(x, plot = FALSE, border=NA,breaks=50)
    breaks <- h$breaks; nB <- length(breaks)
  y <- h$counts; y <- y/max(y)
    rect(breaks[-nB], 0, breaks[-1], y,col=hist.col,border=NA)
}

suppressWarnings(suppressMessages(library(plotly)))
suppressWarnings(suppressMessages(library(reshape)))
''')


for s in samples:
    %R -i s
    display(HTML("<i>Making figures for files from folder " + str(s) + "(" + str(sample
    ro.reval('''
    allqnt <- PSMDat[[s]]

    condNames[[s]] <- ExpDesign[ExpDesign$spec_dir == s,"sample_orig"]
    conditions <- condNames[[s]]
    if (sampledirs[s] != "")
    if(s != "./" & length(sampledirs) > 1) condNames[[s]] <- paste(paste("Sample", 1:le
    pData(allqnt)$sample_name <- paste(condNames[[s]], sampledirs[s])
    pData(allqnt)$sample_group <- condNames[[s]]


    # Better naming:
    samplename <- ifelse(s=="./","",s)

    ## QC Plots
    # violin plots to show distribution of PSM quantifications
    conditionsPlot <- rep(conditions, each=nrow(exprs(allqnt)))
    p <- ggplot(melt(exprs(allqnt)), aes(x=X2, y=value, fill=conditionsPlot)) + geom_vi
    geom_boxplot(width=0.1) + theme_classic() + theme(axis.text.x = element_text(angle
```

```
    xlab("") + ylab("PSM expression") + ggtitle(paste("Distributions in MS run",samplen
    png(filename=paste(sampledirs[s],"/QC_PSM_violinplots.png",sep=""),width=600,height
    print(p)
    dev.off()
    pdf(file=paste(sampledirs[s],"/QC_PSM_violinplots.pdf",sep=""),width=8,height=8)
    print(p)
    dev.off()

    ## histograms of PSMs and peptides per peptide/protein, counting for unique and sha
    QC_PSMHist <- function() {
            par(mfrow=c(2,3))
# all PSMs per peptide
hist(table(fData(allqnt)$Sequence),xlab="PSMs per peptide",100,border=0,col="#AA4444",

    # all PSMs per protein (unique and shared)
AllSingleProts <- gsub(" ","",unlist(strsplit(fData(allqnt)$accession,",")))
hist(table((AllSingleProts)),xlab="PSMs per protein",100,border=0,col="#44AA44", main="

# all PSMS per protein group
hist(table(fData(allqnt)$accession),xlab="PSMs per protein group",100,border=0,col="#44

PeptideCounter <- unique(fData(allqnt)[,c("Sequence","accession")])
# unique and shared peptides per protein
AllSingleProts <- gsub(" ","",unlist(strsplit(PeptideCounter$accession,",")))
hist(table((AllSingleProts)),xlab="Peptides per protein",100,border=0,col="#AAAA44", ma

# all unique peptides per protein group
hist(table(PeptideCounter$accession),xlab="Peptides per protein group",100,border=0,col

# proteins per peptide
hist(sapply(strsplit(PeptideCounter$accession,","), length),xlab="Proteins matching pep
    }
    png(filename=paste(sampledirs[s],"/QC_PSM_and_peptide_distribution.png",sep=""),wid
    QC_PSMHist()
    dev.off()
    pdf(file=paste(sampledirs[s],"/QC_PSM_and_peptide_distribution.pdf",sep=""),width=1
    QC_PSMHist()
    dev.off()

    par(mfrow=c(1,1))
    # Pairwise comparison of samples within on MS run
    QC_Pairs <- function () {
      pairs(exprs(allqnt),lower.panel=panel.smooth, upper.panel=panel.cor, diag.panel=p
        main = paste("MS run",samplename),
        cex=0.1,col="#33333388",pch=15, labels =  condNames[[s]])
    }
    png(filename=paste(sampledirs[s],"/QC_Pairwise_comparison.png",sep=""),width=800,he
    QC_Pairs()
```

```
dev.off()
pdf(file=paste(sampledirs[s],"/QC_Pairwise_comparison.pdf",sep=""),width=15,height=
QC_Pairs()
dev.off()

''')
# display figures

%R --width 800  print(p);QC_PSMHist();par(mfrow=c(1,1));QC_Pairs()

ro.reval('''
PSMDat[[s]] <- allqnt
''')
```

```
<IPython.core.display.HTML object>
```

```
<IPython.core.display.HTML object>
```

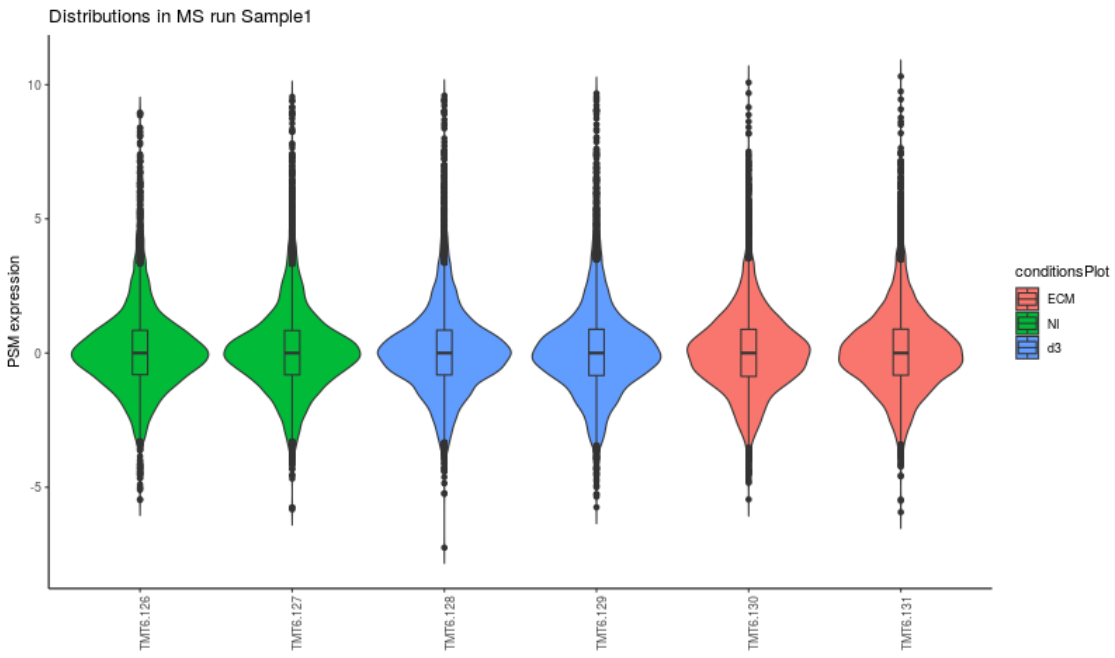

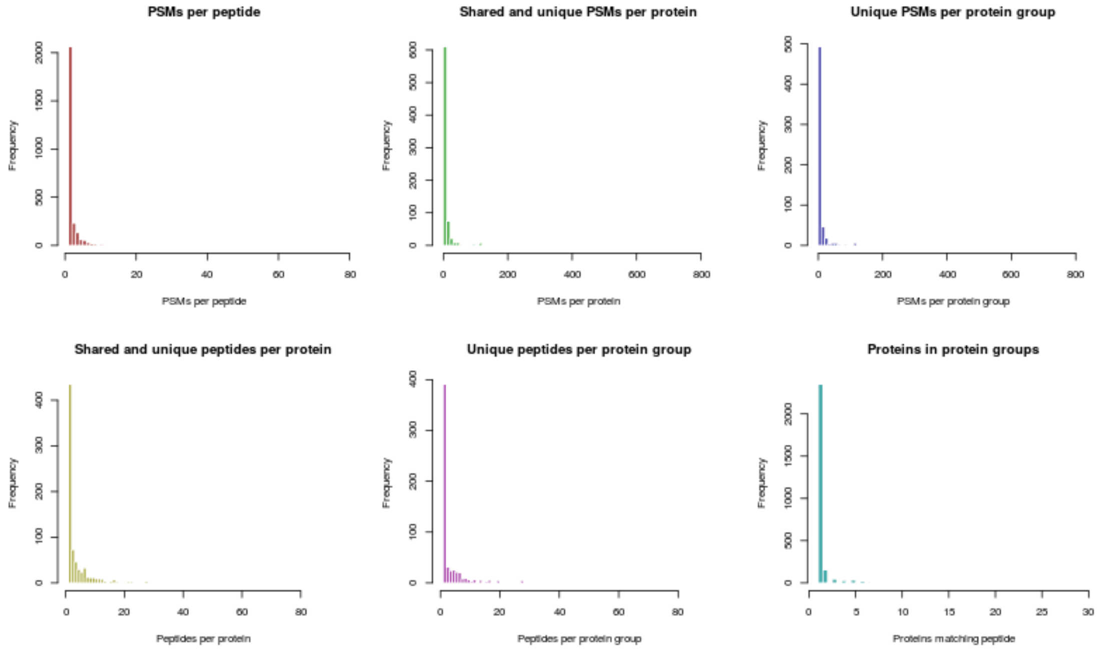


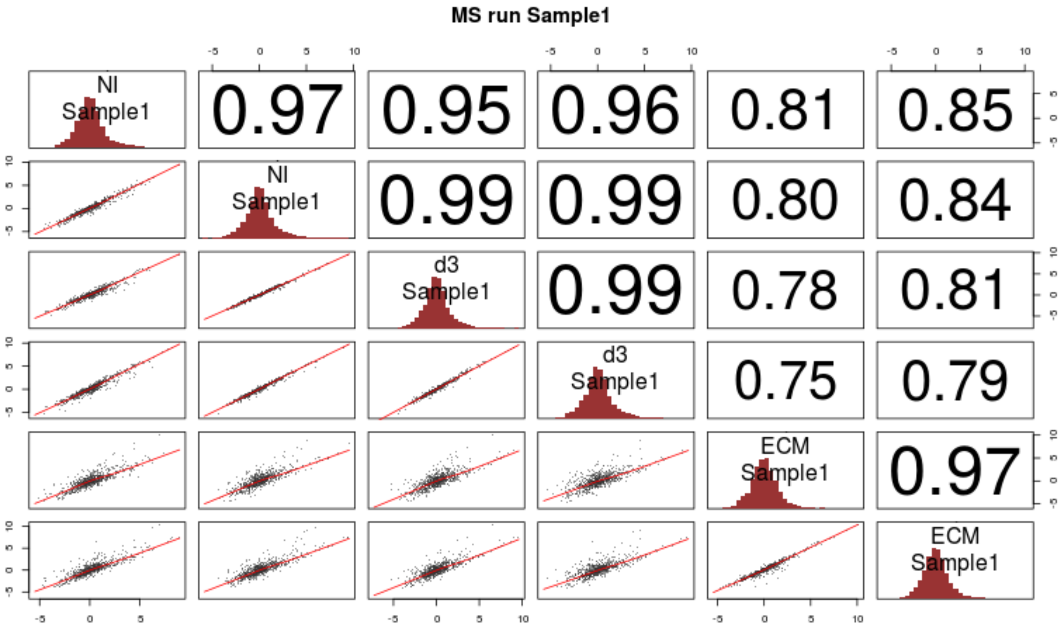


<IPython.core.display.HTML object>

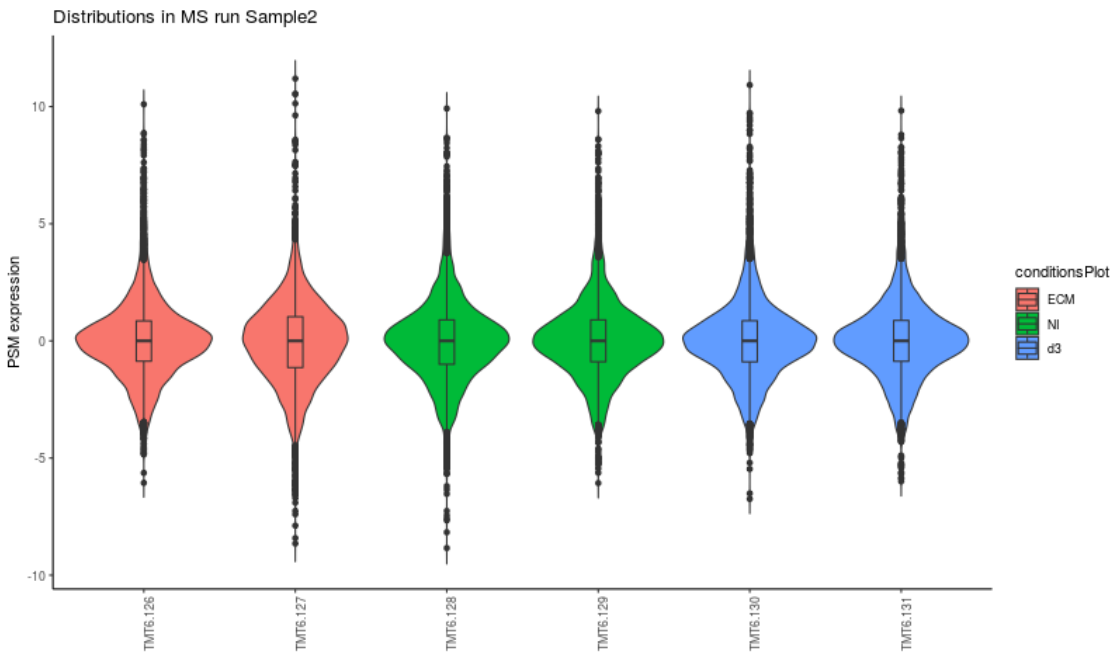
Distributions in MS run Sample2
PSM expression
TMT6.126
TMT6.127
TMT6.128
TMT6.129
TMT6.130
TMT6.131
conditionsPlot
ECM
NI
d3

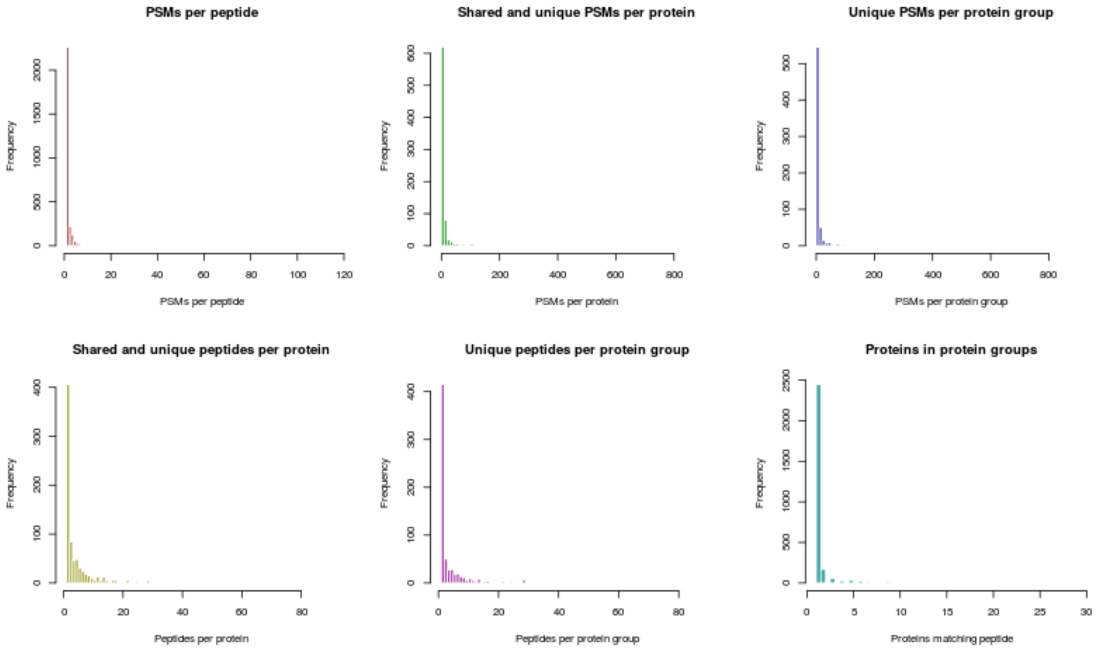
PSMs per peptide
Shared and unique PSMs per protein
Unique PSMs per protein group
Shared and unique peptides per protein
Unique peptides per protein group
Proteins in protein groups
Frequency
PSMs per peptide
PSMs per protein
PSMs per protein group
Peptides per protein
Peptides per protein group
Proteins matching peptide

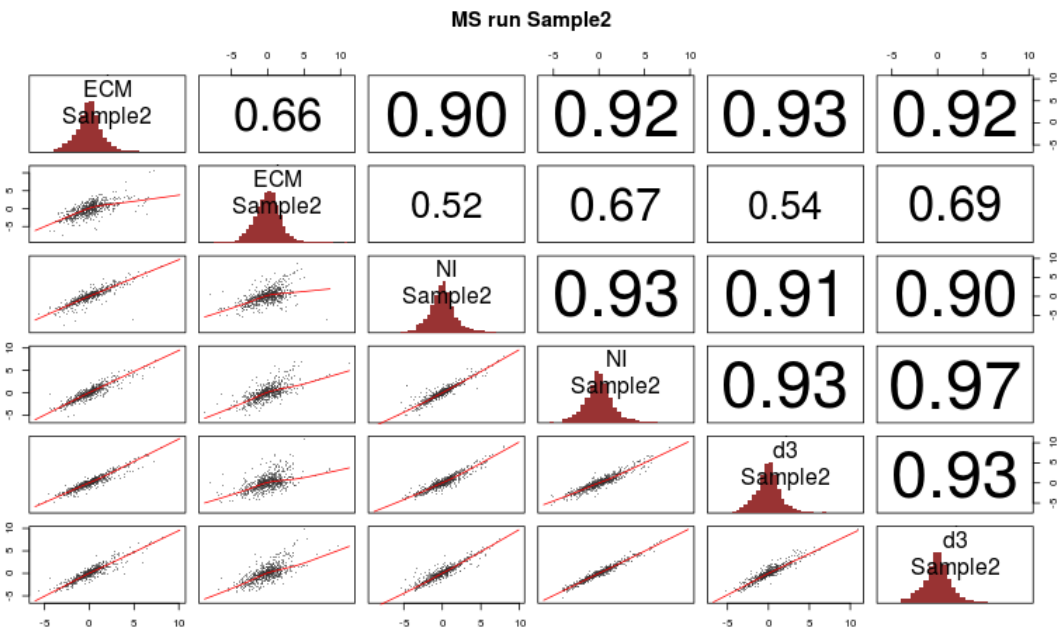


#### 2.1.4 Protein Inference

```
In [52]: display(HTML("<h4>Protein quantification ...</h4>"))

         ro.reval('''
         summarization_function <- search_in["summarization_method"][[1]]

         # get minimum peptides
         min_peps <- search_in["min_protein_psms"][[1]]
         ''')

         display(HTML("Using " + str(ro.r.summarization_function[0]) + " for protein summarizati
         display(HTML("and requiring a minimum of " + str(ro.r.min_peps[0]) + " PSMs per protein

         for s in samples:
             %R -i s
             display(HTML("<i>Making figures for files from folder " + str(s) + " (" + str(sampl
             display(HTML("Filtering and summarization"))
             ro.reval('''
             load(paste(sampledirs[s],"/AllQuantPSMs.RData",sep=""))

             allqnt <- PSMDat[[s]]

             ### TODO: remove only previously specified variable PTMs
             # Remove other PTMs for quantification
```

```
<IPython.core.display.HTML object>

<IPython.core.display.HTML object>

<IPython.core.display.HTML object>

<IPython.core.display.HTML object>

<IPython.core.display.HTML object>
```

```
Removing 3 PSMs with more than 50% missing values for quantification

Removing 1570 PSMs corresponding to 499 proteins with less then 2 peptides

Your data contains missing values. Please read the relevant section in
the combineFeatures manual page for details the effects of missing
values on data aggregation.
```

```
<IPython.core.display.HTML object>
```

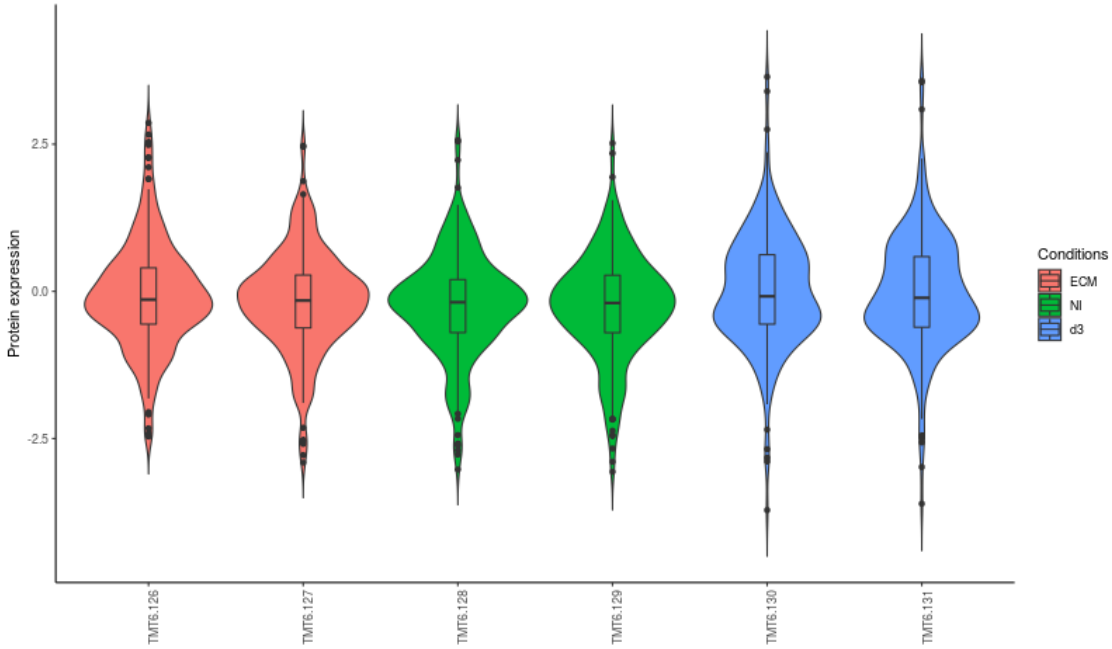


```
<IPython.core.display.HTML object>
```

```
<IPython.core.display.HTML object>
```

```
Removing 2 PSMs with more than 50% missing values for quantification

Removing 1816 PSMs corresponding to 570 proteins with less then 2 peptides

Your data contains missing values. Please read the relevant section in
the combineFeatures manual page for details the effects of missing
values on data aggregation.
```

```
<IPython.core.display.HTML object>
```

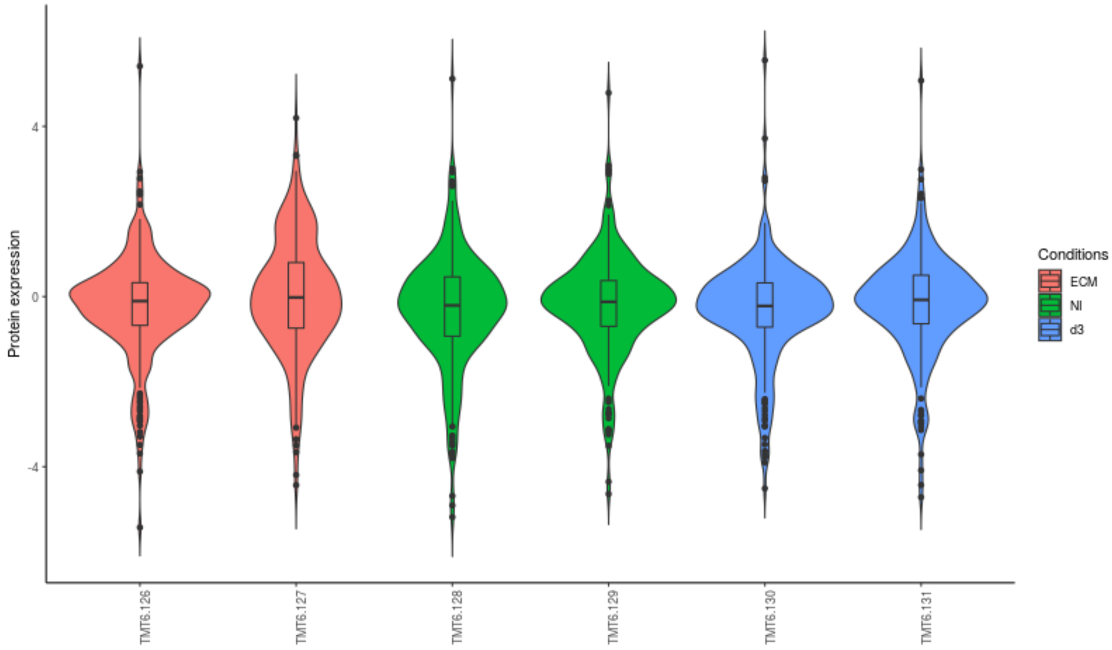


#### 2.1.5 Sample similarity and statistics

In [53]:
```
display(HTML("<h4>Protein quantification ...</h4>"))

display(HTML("Merging sample runs from different folders ..."))
ro.reval('''
par(mfrow=c(1,1))

# Adjust each protein of each MS run by the mean of their expressions over the channels
# or reference condition (e.g. pool)?

for (s in samples) {
    norm_vec <- 0
    if (search_in["ref_condition"] == "Mean over channels") {
        norm_vec <- rowMeans(ProtDat[[s]],na.rm=T)
    } else {
        ref_cols <- which(ExpDesign$sample_group[ExpDesign$spec_dir == s] == search_in[
        if (length(ref_cols) > 0) {
            norm_vec <- rowMeans(ProtDat[[s]][, ref_cols, drop=F],na.rm=T)
        } else {
            message(paste("No reference samples in MS run",s,"! Taking mean of all chan
            norm_vec <- rowMeans(ProtDat[[s]],na.rm=T)
        }
```

```
<IPython.core.display.HTML object>
```

<IPython.core.display.HTML object>

[1]
 "Merging samples (if needed) ..."

<IPython.core.display.HTML object>

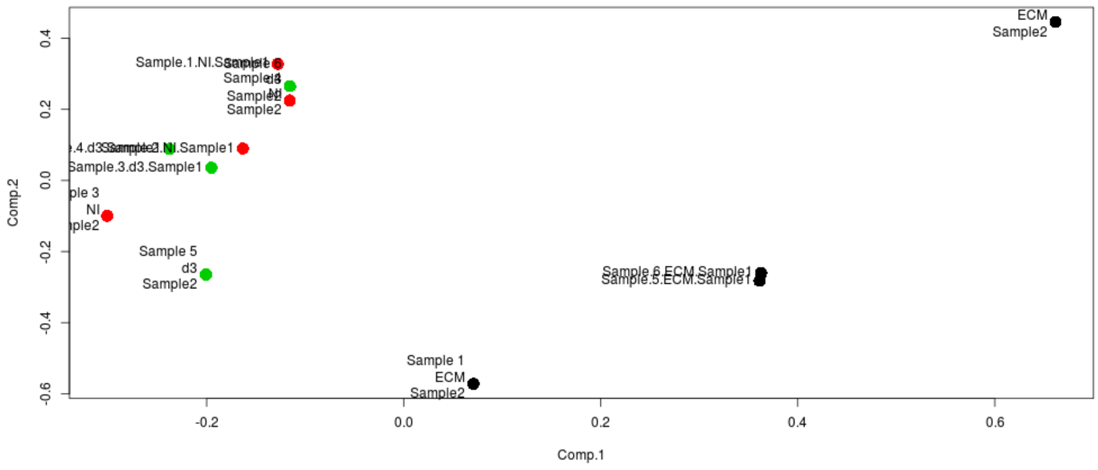


<IPython.core.display.HTML object>

<IPython.core.display.HTML object>

[1]
 "Statistical tests carried out to compare: d3-NI"

[2]
 "Statistical tests carried out to compare: ECM-NI"

<IPython.core.display.HTML object>

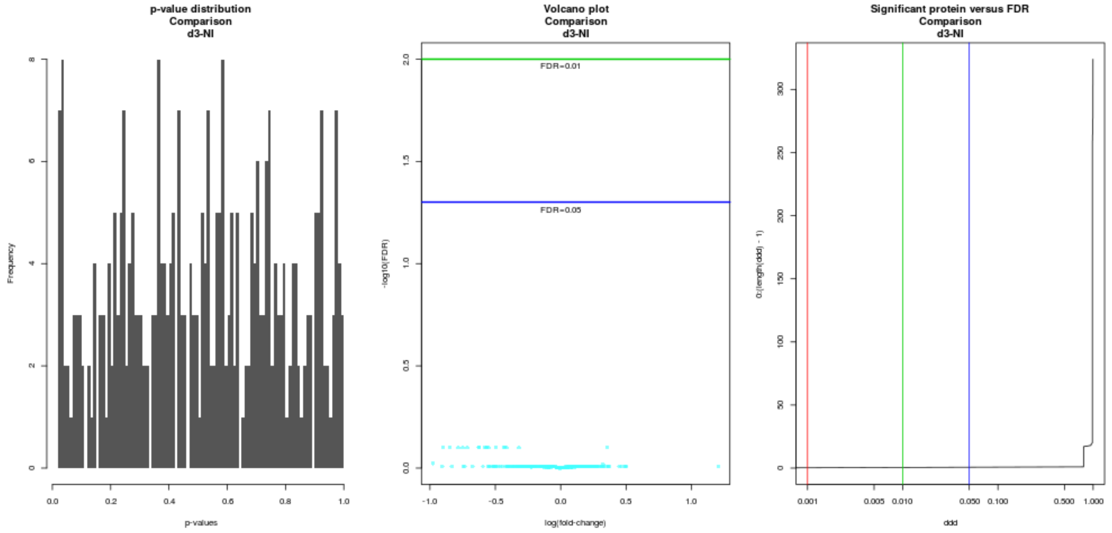


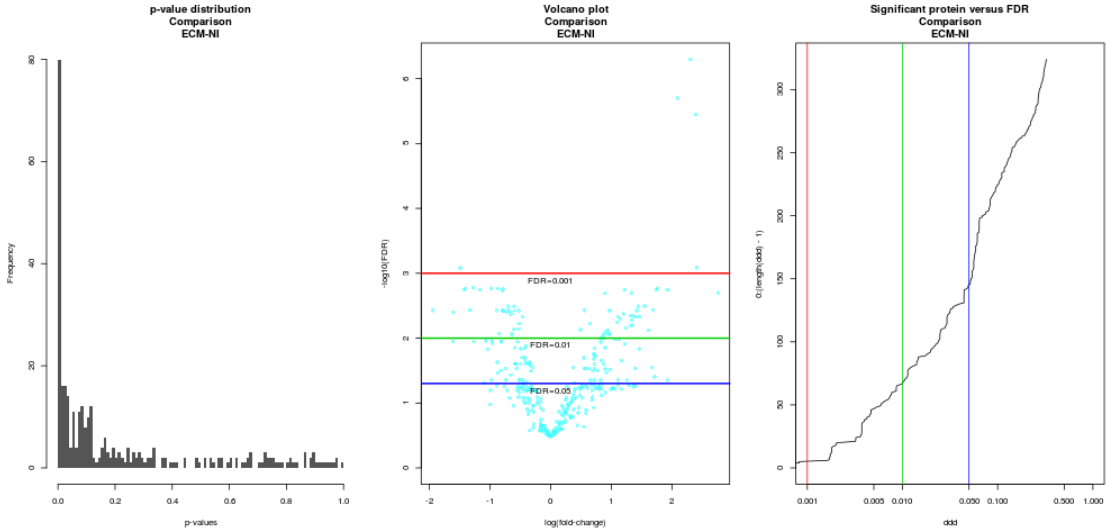


#### 2.1.6 Run your own script
