## Supplementary File 3 for "IsoProt: A complete and reproducible workflow to analyse iTRAQ/TMT experiments"

```
Tab(children=(VBox(children=(HTML(value='<h4>Folders and files</h4>'), HTML(value='Folder for sp
```

```
<IPython.core.display.Javascript object>
```

```
ExecuteButton(button_text='Run R analysis', disabled=True, n_next_cells=13, style=ButtonStyle())
```

```
Saving parameters to /home/biodocker/OUT/protocol_parameters.json
Deleting existing files in /home/biodocker/OUT
Adapting MGF titles...
['/home/biodocker/data/iTRAQ_fractionation_2.mgf', '/home/biodocker/data/iTRAQ_fractionation_1-4
Extracting reporter peaks...
Creating decoy database...
Start: Creating decoy database...
    Completed in 2 sec
Start: Creating search parameter file
    Completed in 0 sec
Start: SearchCLI
    Completed in 42:7 min
Start: PeptideShakerCLI processing
    Completed in 3:17 min
```

```
Start: ReportCLI (conversion to .tsv)
    Completed in 1:59 min
Search Done.
Start the analysis by clicking the >>Run R Analysis<< button
Saving parameters to /home/biodocker/OUT/protocol_parameters.json
Saving parameters to /home/biodocker/OUT/protocol_parameters.json
Deleting existing files in /home/biodocker/OUT
Adapting MGF titles...
['/home/biodocker/data/iTRAQ_fractionation_2.mgf', '/home/biodocker/data/iTRAQ_fractionation_1-4
Extracting reporter peaks...
Creating decoy database...
Start: Creating decoy database...
    Completed in 2 sec
Start: Creating search parameter file
    Completed in 0 sec
Start: SearchCLI
    Completed in 44:7 min
Start: PeptideShakerCLI processing
    Completed in 3:19 min
Start: ReportCLI (conversion to .tsv)
    Completed in 2:7 min
Search Done.
Start the analysis by clicking the >>Run R Analysis<< button
Saving parameters to /home/biodocker/OUT/protocol_parameters.json
Saving parameters to /home/biodocker/OUT/protocol_parameters.json
```

### 2.1 Methods Sections

```
In [19]: %run Scripts/methods_generator.ipy
```

```
HTML(value="<b>Citations are added as PubMed ids</b><br />The mass spectrometry data was analyse
```

#### 2.1.1 Convert Search Settings into R Objects

These cells should not produce any output.

```
In [20]: # Since py2ri cannot convert dict objects, simply save everything R needs as a JSON str

         # remove the callback function first
         search_in.pop("on_search_complete", None)

         import json
         search_in_string = json.dumps(search_in)
         search_out_string = json.dumps(search_out)

         import rpy2.robjects as ro
```

```python
from IPython.display import display,HTML
import sys
```

In [21]:
```r
%%R -i search_in_string,search_out_string -o samples
# Convert the JSON objects back into "natural" R objects
suppressWarnings(library(rjson))

search_in = fromJSON(search_in_string, simplify = T)
search_out = fromJSON(search_out_string, simplify = T)

# load the experimental design
ExpDesign <- read.table(search_in["exp_design_file"][[1]],sep="\t",header=T)

rm(search_in_string, search_out_string)


## assuming that the computer allows min. 4 threads
NumThreads <- 4

# library causes the execution to fail if the library is missing
suppressWarnings(suppressMessages(library(lattice)))
suppressWarnings(suppressMessages(library(stringr)))
suppressWarnings(suppressMessages(library(mzID)))
suppressWarnings(suppressMessages(library(MSnbase)))

# warnings as stdout
sink(stdout(), type = "message")

message("Verifying availability of files ...")

# change to the input directory
spectradirs <- unlist(search_in["input_directory"])
out_dir <- search_in["workdir"][[1]]
#setwd(out_dir)

## Folder names for different runs (e.g. different replicates, ...)
samples <- unique(as.character(ExpDesign$spec_dir))

# If only one folder, then set correct folder names
if(length(samples) == 1) {
    samples <- "./"
    ExpDesign$spec_dir <- "./"
}
sampledirs <- paste(out_dir,"/",samples,sep="")
all_ident_files <- search_out[[1]]
if (length(sampledirs) != length(spectradirs) | length(sampledirs) != length(all_ident_
    stop("Unequal number of sample folders")
}
```

```
        names(sampledirs) <- names(spectradirs) <- names(all_ident_files) <- samples

        message("OK")
```

```
Verifying availability of files ...
OK
```

#### 2.1.2 Load and Quantify Spectra at the peptide level

```
In [22]: display(HTML("<h4>Processing identification data...</h4>"))

         def loadPsms(s):
             """
             Loads the PSMs of the specified samples in the R environment.
             :param s: The sample to be processed
             :return: Nothing is returned. A "psms" object is created in the R environment.
             """
             %R -i s

             ro.reval('''
             # warnings as stdout
             sink(stdout(), type = "message")

             ident_files <- all_ident_files[s]

             if (is.null(ident_files)) stop("Error: No identification files")

             if (!file.exists(ident_files)) stop(paste("Error: Cannot find identification files"

             # ---- load the PSMs ----
             max_fdr <- search_in["target_fdr"]
             psms <- read.csv(ident_files, sep = "\t",stringsAsFactors = F)

             if (! "Decoy" %in% names(psms)) {
                 stop("Error: No decoy information available in output file")
             }

             num_psms <- nrow(psms)
             ''')

             num_psms = ro.r.num_psms[0]
             display(HTML(" - Loaded " + str(num_psms) + " PSMs"))

             ro.reval('''
             # ---- Confidence filter ----
             # TODO: This could be replaced by the PeptideShaker functionality
```

```
<IPython.core.display.HTML object>
```

```
<IPython.core.display.HTML object>
```

```
<IPython.core.display.HTML object>
```

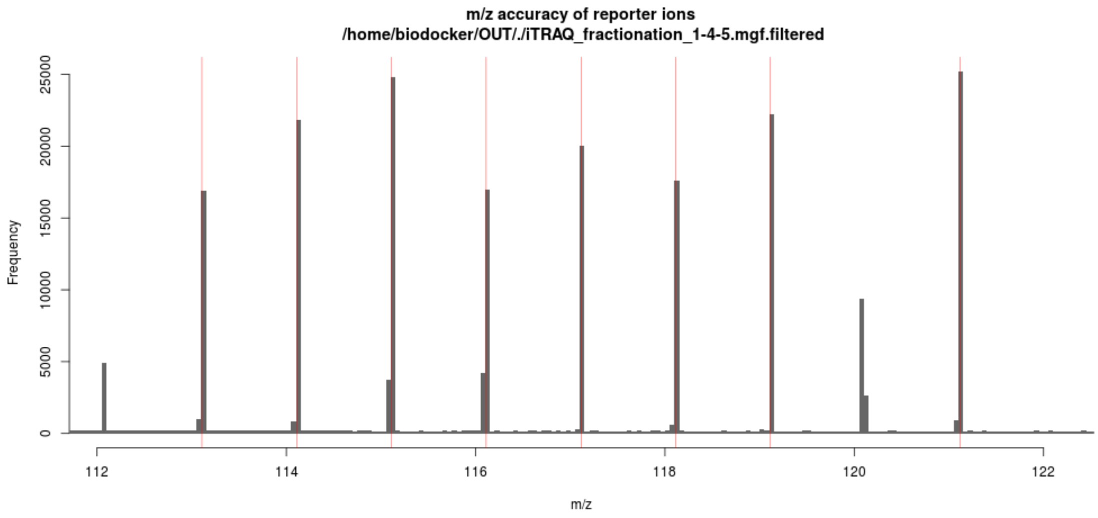

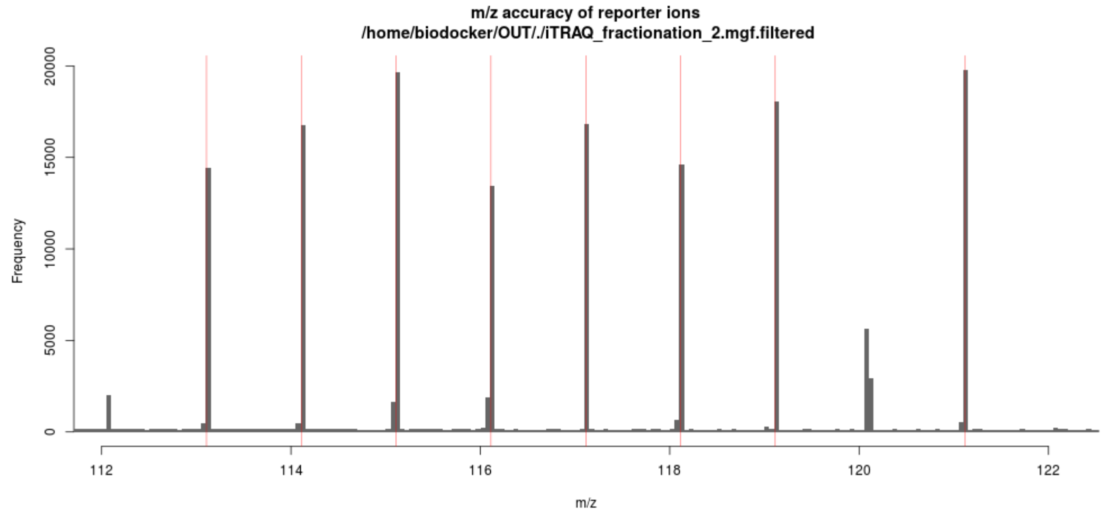


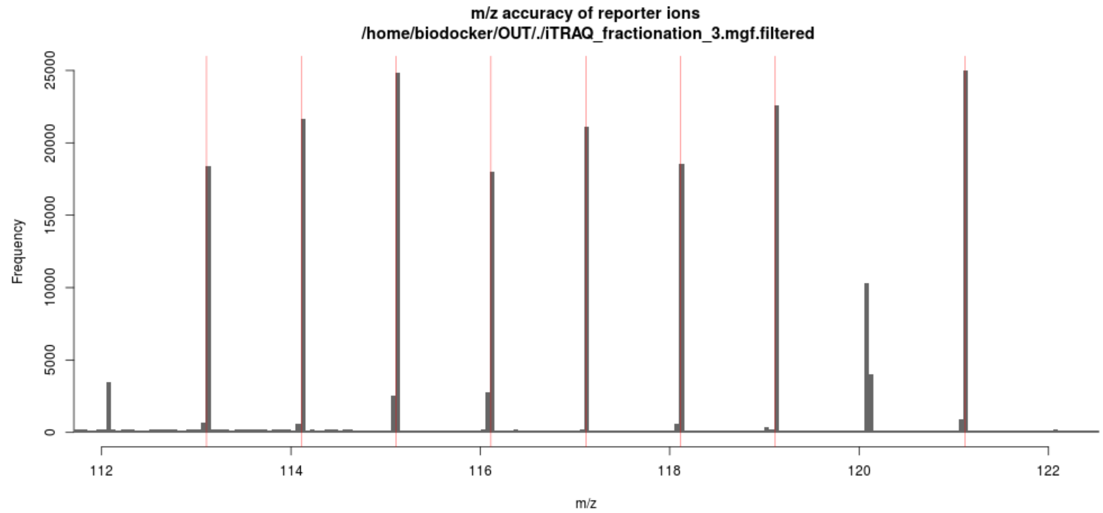


```
<IPython.core.display.HTML object>
```

```
  Removed 15620 features with 50% missing values

  Removed 13130 features with 50% missing values

  Removed 11192 features with 50% missing values
```

```
<IPython.core.display.HTML object>
```

```
<IPython.core.display.HTML object>
```

```
<IPython.core.display.HTML object>
```

#### 2.1.3 Basic Quality Control

```
In [23]: display(HTML("<h4>Processing PSMs ...</h4>"))

         %R condNames  <- list()

         ro.reval('''
         # Used panel functions
         panel.cor <- function(x, y, digits=2, prefix="", cex.cor, ...)
         {
             usr <- par("usr"); on.exit(par(usr))
             par(usr = c(0, 1, 0, 1))
             r <- abs(cor(x, y, use="na.or.complete"))
             txt <- format(c(r, 0.123456789), digits=digits)[1]
             txt <- paste(prefix, txt, sep="")
             cex.cor <- 0.8/strwidth(txt)
             test <- cor.test(x,y)
             # borrowed from printCoefmat
             Signif <- symnum(test$p.value, corr = FALSE, na = FALSE,
                           cutpoints = c(0, 0.001, 0.01, 0.05, 0.1, 1),
                           symbols = c("***", "**", "*", ".", " "))

             text(0.5, 0.5, txt, cex = cex.cor * r)
         }

         panel.hist <- function(x, hist.col="#993333", ...)
         {
             usr <- par("usr"); on.exit(par(usr))
            par(usr = c(usr[1:2], 0, 1.5) )
             h <- hist(x, plot = FALSE, border=NA,breaks=50)
             breaks <- h$breaks; nB <- length(breaks)
           y <- h$counts; y <- y/max(y)
             rect(breaks[-nB], 0, breaks[-1], y,col=hist.col,border=NA)
         }

         suppressWarnings(suppressMessages(library(plotly)))
         suppressWarnings(suppressMessages(library(reshape)))
         ''')
```

```
<IPython.core.display.HTML object>

<IPython.core.display.HTML object>
```

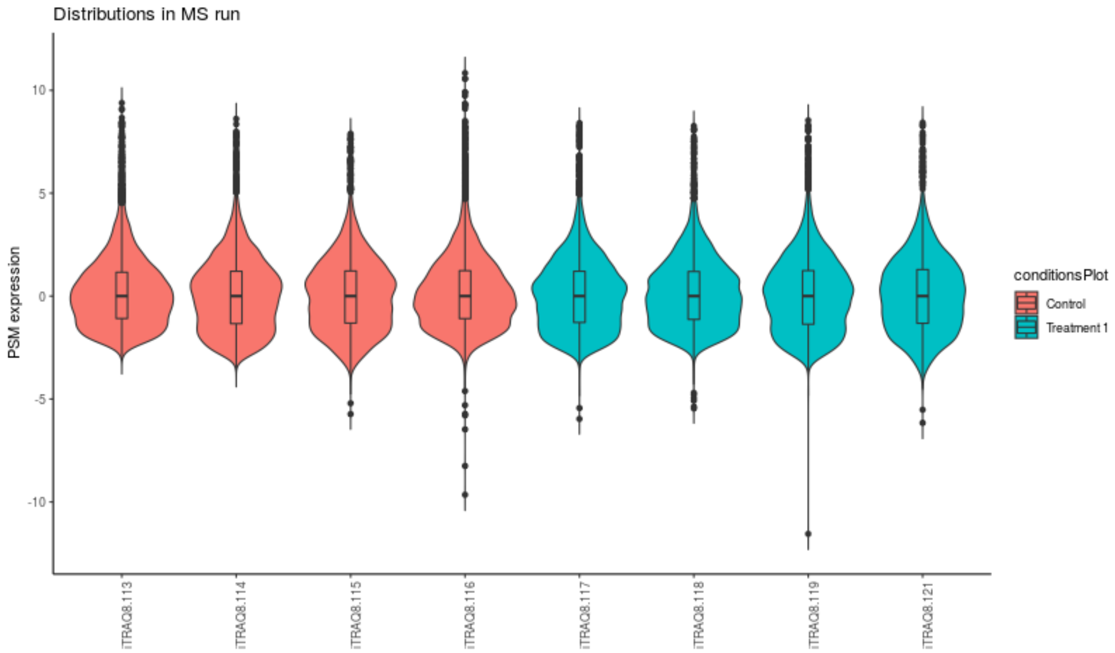
Distributions in MS run
PSM expression
10
5
0
-5
-10
iTRAQ8.113
iTRAQ8.114
iTRAQ8.115
iTRAQ8.116
iTRAQ8.117
iTRAQ8.118
iTRAQ8.119
iTRAQ8.121
conditionsPlot
Control
Treatment 1

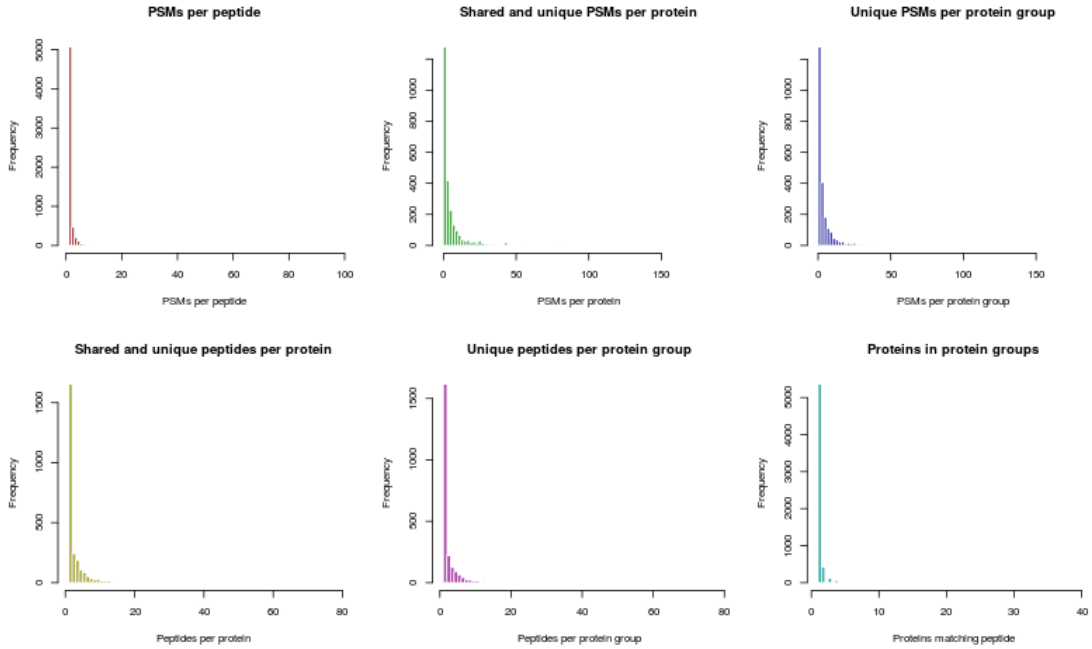
PSMs per peptide
Frequency
PSMs per peptide
Shared and unique PSMs per protein
Frequency
PSMs per protein
Unique PSMs per protein group
Frequency
PSMs per protein group
Shared and unique peptides per protein
Frequency
Peptides per protein
Unique peptides per protein group
Frequency
Peptides per protein group
Proteins in protein groups
Frequency
Proteins matching peptide

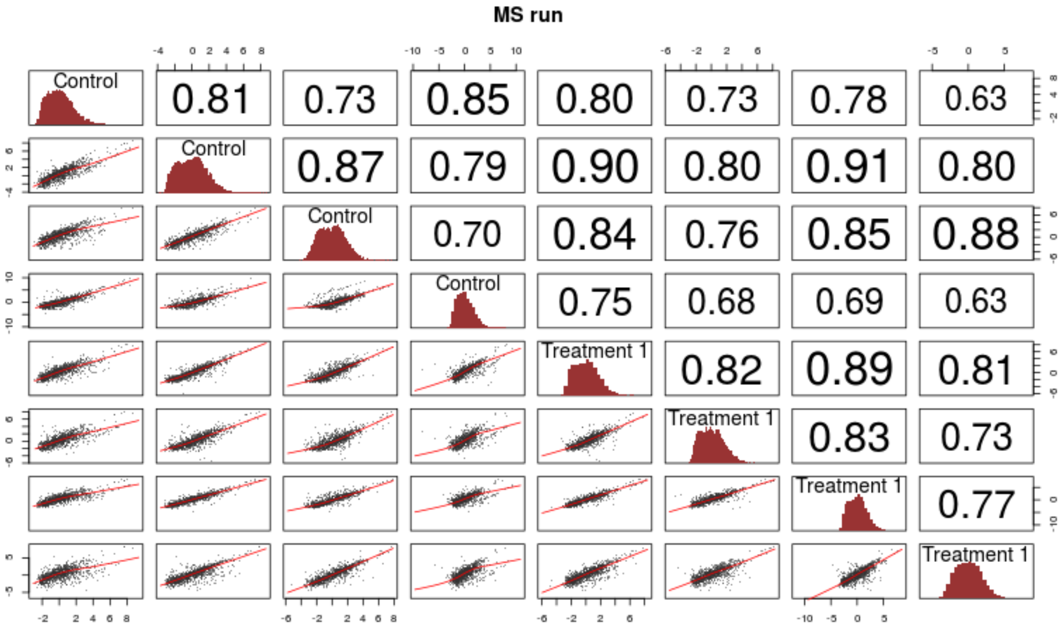


#### 2.1.4 Protein Inference

```
In [24]: display(HTML("<h4>Protein quantification ...</h4>"))

         ro.reval('''
         summarization_function <- search_in["summarization_method"][[1]]

         # get minimum peptides
         min_peps <- search_in["min_protein_psms"][[1]]
         ''')

         display(HTML("Using " + str(ro.r.summarization_function[0]) + " for protein summarizati
         display(HTML("and requiring a minimum of " + str(ro.r.min_peps[0]) + " PSMs per protein

         for s in samples:
             %R -i s
             display(HTML("<i>Making figures for files from folder " + str(s) + " (" + str(sampl
             display(HTML("Filtering and summarization"))
             ro.reval('''
             load(paste(sampledirs[s],"/AllQuantPSMs.RData",sep=""))

             allqnt <- PSMDat[[s]]

             ### TODO: remove only previously specified variable PTMs
             # Remove other PTMs for quantification
```

```
<IPython.core.display.HTML object>


<IPython.core.display.HTML object>


<IPython.core.display.HTML object>


<IPython.core.display.HTML object>


<IPython.core.display.HTML object>
```

```
Removing 11 PSMs with more than 50% missing values for quantification

Removing 3951 PSMs corresponding to 1864 proteins with less then 2 peptides

Your data contains missing values. Please read the relevant section in
the combineFeatures manual page for details the effects of missing
values on data aggregation.



<IPython.core.display.HTML object>
```

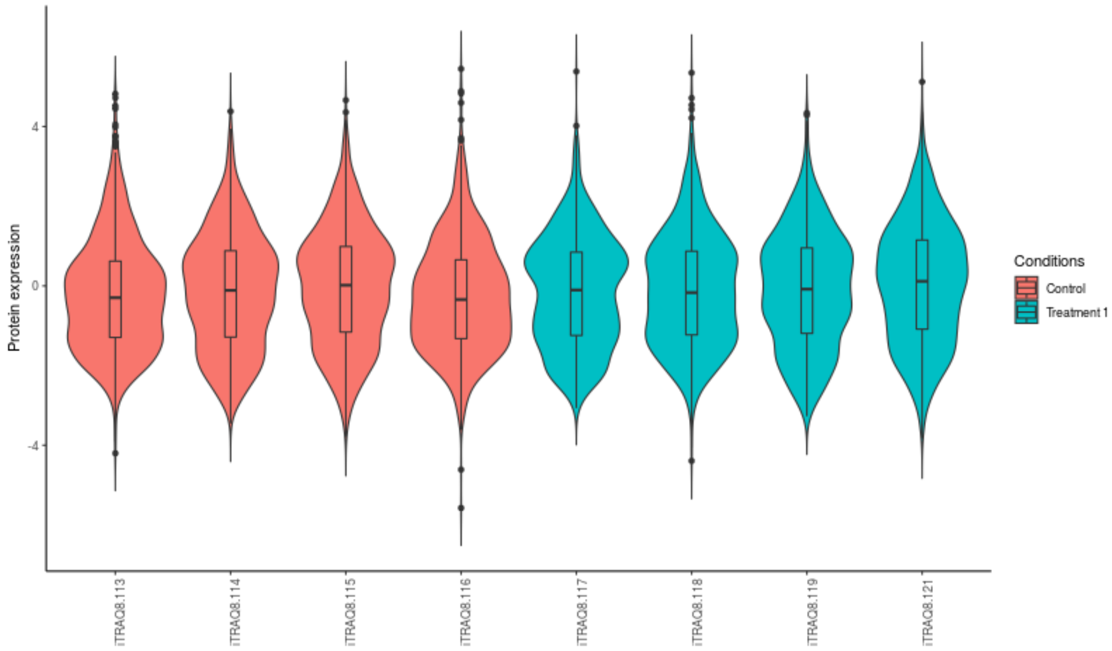


##### 2.1.5 Sample similarity and statistics

```
In [25]: display(HTML("<h4>Protein quantification ...</h4>"))

         display(HTML("Merging sample runs from different folders ..."))
         ro.reval('''
         par(mfrow=c(1,1))

         # Adjust each protein of each MS run by the mean of their expressions over the channels
         # or reference condition (e.g. pool)?

         for (s in samples) {
             norm_vec <- 0
```

```
    if (search_in["ref_condition"] == "Mean over channels") {
        norm_vec <- rowMeans(ProtDat[[s]],na.rm=T)
    } else {
        ref_cols <- which(ExpDesign$sample_group[ExpDesign$spec_dir == s] == search_in[
        if (length(ref_cols) > 0) {
            norm_vec <- rowMeans(ProtDat[[s]][, ref_cols, drop=F],na.rm=T)
        } else {
            message(paste("No reference samples in MS run",s,"! Taking mean of all chan
            norm_vec <- rowMeans(ProtDat[[s]],na.rm=T)
        }
    }
    ProtDat[[s]] <- ProtDat[[s]] - norm_vec
}


# Merge different samples
    totProts <- ProtDat[[1]]
if (length(samples)>1) {
    print("Merging samples (if needed) ...")
    totProts <- data.frame(rownames(totProts),totProts)
    for (s in samples[2:length(samples)])
        totProts <- merge(totProts,ProtDat[[s]], all=T, by.x=1, by.y=0)
    rownames(totProts) <- totProts[,1]
    totProts <- totProts[,2:ncol(totProts)]
}
write.csv(totProts,file=paste(out_dir,"/AllQuantProteinsInAllSamples.csv",sep=""))
''')

display(HTML("Principal component analysis ..."))
ro.reval('''
# PCA
pca <- princomp((totProts[complete.cases((totProts)),]))
  #plot(pca)

QC_PCA <- function() {
    plot(pca$loadings, cex=2, pch=16, col=as.numeric(as.factor(ExpDesign[["sample_orig"
    text(pca$loadings,colnames(totProts), pos=2)
}
png(filename=paste(out_dir,"/QC_Stat_PCA.png",sep=""),width=800,height=800)
QC_PCA()
dev.off()
pdf(file=paste(out_dir,"/QC_Stat_PCA.pdf",sep=""),width=8,height=8)
QC_PCA()
dev.off()
num_prots <- nrow(allProts)
''')
# display figure
%R --width 1000 QC_PCA()
```

```
display(HTML("Quantified a total of " + str(ro.r.num_prots[0]) + " protein groups"))

display(HTML("Running LIMMA for statistical tests ..."))

ro.reval('''
##Statistics
library(limma)
library(qvalue)
NumCond <- length(unique(ExpDesign$sample_orig))
  if (NumCond < 2)
      stop("Only 1 experimental condition -> no statistics")

design <- model.matrix(~0+factor(ExpDesign$sample_group)-1)
  colnames(design)<-make.names(paste(unique(ExpDesign$sample_orig),sep=""))
  contrasts<-NULL
  First <- which(unique(ExpDesign$sample_group) == search_in[["stat_condition"]])
  for (i in (1:NumCond)[-First]) contrasts<-append(contrasts,paste(colnames(design)[i],
  print(paste("Statistical tests carried out to compare:",contrasts))
  contrast.matrix<-makeContrasts(contrasts=contrasts,levels=design)
  # print(dim(Data))
  lm.fitted <- lmFit(totProts,design)
  lm.contr <- contrasts.fit(lm.fitted,contrast.matrix)
  lm.bayes <- eBayes(lm.contr)
  #topTable(lm.bayes)
 plvalues <- lm.bayes$p.value
fcs <- lm.bayes$coefficients
  qlvalues <- matrix(NA,nrow=nrow(plvalues),ncol=ncol(plvalues),dimnames=dimnames(plval
  # qvalue correction
 for (i in 1:ncol(plvalues)) {
    tqs <- qvalue(na.omit(plvalues[,i]))$qvalues
    qlvalues[names(tqs),i] <- tqs
  }


par(mfrow=c(1,3))

statTable <- NULL

# Visualizations: volcano plot, number of regulated proteins per FDR, interactive table

# p-value distributions

QC_pvals <- function(i) {
    hist(plvalues[,i],100,border=NA,col="#555555", main=paste("p-value distribution\nCo
        xlab="p-values")
}
# volcano plots
```

```
QC_volcanos <- function(i) {
    plot(fcs[,i], -log10(qlvalues[,i]),pch=16,col="#44FFFF99",
        xlab="log(fold-change)", ylab="-log10(FDR)", main=paste("Volcano plot\nComparis
                                                    colnames(fcs)[i], sep="\n")
        ylim=c(0,max(2,max(-log10(qlvalues[,i]), na.rm=T))))
    abline(h=-log10(c(0.001,0.01,0.05)), col=2:4,lwd=2)
    text(c(0,0,0), c(3,2,1.3),c("FDR=0.001","FDR=0.01","FDR=0.05"),pos=1)
}
# Number of sigificant features vs. FDR
QC_NumSig <- function(i) {
    ddd <- c(0.0001,sort(qlvalues[,i]))
    plot(ddd, 0:(length(ddd)-1),type="l",xlim=c(1e-3,1),log="x", main=paste("Significan
    abline(v=c(0.001,0.01,0.05), col=2:4, xlab="FDR", ylab="Number of proteins")
}
QC_AllStats <- function(i) {
    QC_pvals(i)
    QC_volcanos(i)
    QC_NumSig(i)
}

for (i in 1:ncol(plvalues)) {
    # export figures to pdf/png
    png(filename=paste(out_dir,"/QC_Stat_Summary_",colnames(fcs)[i],".png",sep=""),widt
    par(mfrow=c(1,3))
    QC_pvals(i)
    QC_volcanos(i)
    QC_NumSig(i)
    dev.off()
    pdf(file=paste(out_dir,"/QC_Stat_Summary_",colnames(fcs)[i],".pdf",sep=""),width=15
    par(mfrow=c(1,3))
    QC_pvals(i)
    QC_volcanos(i)
    QC_NumSig(i)
    dev.off()
par(mfrow=c(1,1))

}
''')

display(HTML("Saving statistics results ..."))
ro.reval('''
# How far should we go? Clustering (when having more then 2 groups)?
# Hierarchical clustering of significant features? clusterProfiler?


# Saving stats
statOut <- cbind(fcs, plvalues, qlvalues)
colnames(statOut) <- paste(rep(c("log(fold-change)","p-values","q-values"), each=ncol(p
```

```
statOut <- cbind(Proteins=rownames(fcs),statOut)
#print(head(statOut))
write.csv(statOut, paste(out_dir,"/DifferentiallyRegulatedProteins.csv",sep=""),row.nam

''')

%R -w 1000  aa<-c(1,3); bb <- ncol(plvalues); par(mfrow=aa); for (i in 1:bb) QC_AllStat
```

<IPython.core.display.HTML object>

<IPython.core.display.HTML object>

<IPython.core.display.HTML object>

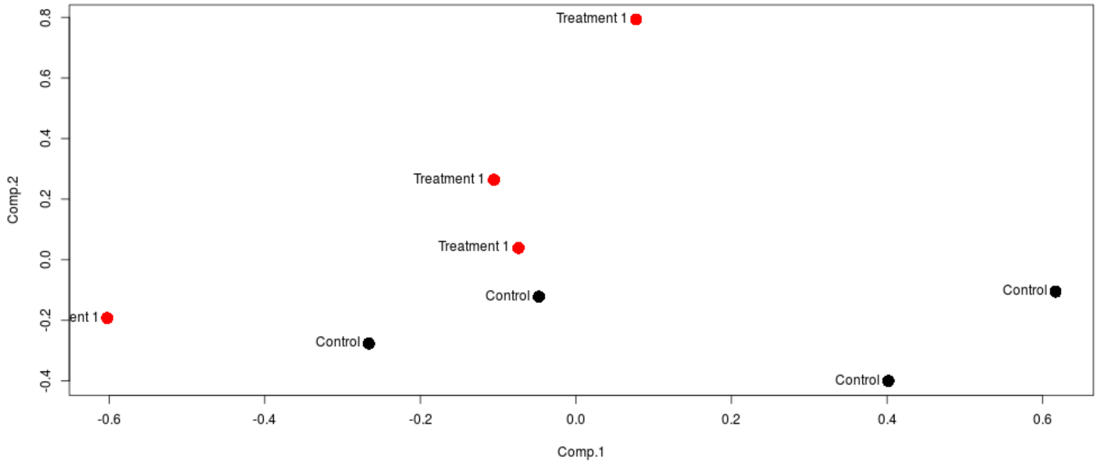


<IPython.core.display.HTML object>

<IPython.core.display.HTML object>

[1]
 "Statistical tests carried out to compare: Treatment.1-Control"

<IPython.core.display.HTML object>

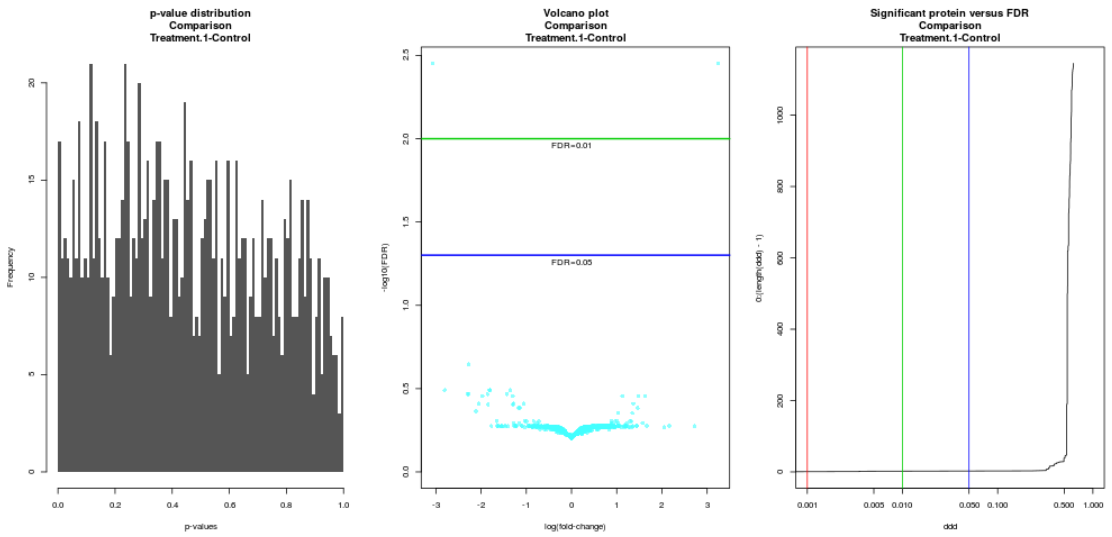


#### 2.1.6 Run your own script

In [ ]:
